# Spike inference from calcium imaging data acquired with GCaMP8 indicators

**DOI:** 10.1101/2025.03.03.641129

**Authors:** Peter Rupprecht, Márton Rózsa, Xusheng Fang, Karel Svoboda, Fritjof Helmchen

## Abstract

For neuroscience to be reproducible, its key methods must be quantitative and interpretable. Calcium imaging is such a key method, but it records neuronal activity only indirectly and is therefore difficult to interpret. These difficulties arise from the kinetics, nonlinearity, and sensitivity of calcium indicators, but also from the methods used for signal analysis. Here, we evaluate how methods for spike inference can be optimized to interpret data with the recently developed calcium indicator GCaMP8. We find that the indicator linearity of GCaMP8 enables more accurate recovery of high-frequency spiking events and the detection of single action potentials. Ground truth recordings from mouse neocortex show that the most linear calcium variants, GCaMP8s and GCaMP8m – but not GCaMP6, GCaMP7f, or GCaMP8f – robustly detect isolated spikes in pyramidal neurons under realistic noise conditions. In addition, we fine-tune and benchmark existing algorithms for spike inference (CASCADE, OASIS, MLSpike) with GCaMP8 data, investigate spike inference for interneurons, and demonstrate how the fast rise time of GCaMP8 enables low-latency real-time detection of neuronal activity. Together, our study provides tools and guidelines to optimally process calcium signals with GCaMP8 and highlights the key role of linearity in improving the interpretability of calcium imaging.

## Introduction

Over the past decades, systems neuroscience has advanced at a rapid pace. This progress has been driven by the astonishingly fast development of new tools for circuit neuroscience, such as large-scale recording techniques and optogenetics^1–3^. Calcium imaging is one of the most powerful examples of these tools, ideally suited to study neural activity at the levels of single cells and across large populations with sub-second temporal resolution^4^. The functional basis of calcium imaging to measure neuronal activity is the influx of calcium ions during action potentials (“spikes”)^5^. The resulting changes of the intracellular calcium concentration are reflected by the fluorescence changes of the cytosolic calcium-sensitive molecule, the calcium indicator. Often, the baseline-normalized fluorescence changes (ΔF/F) are used as readout of neuronal activity. However, this readout is a noisy, indirect, relatively slow, and often nonlinear proxy of neuronal spiking activity^6–9^. To estimate the true spike patterns from this indirect proxy and to simultaneously denoise the recording, spike inference methods are essential tools to process calcium imaging data^10–14^. Existing methods for spike inference have been optimized for specific calcium indicators, either by tuning the parameters of a model-based approach^10,12,15,16^, or by training supervised methods to fit calibration (ground truth) data^14,17,18^. It is often not clear, though, how these methods can be optimally applied to new calcium indicators.

Early calcium indicators were developed as small fluorescent organic dyes^19^, followed by genetically encoded versions created through protein engineering^20,21^. Over the last two decades, the field has largely shifted to genetically encoded calcium indicators^9^, which are continually refined in order to reflect neuronal activity more closely in their fluorescence signals^22–25^. For the most widely used GCaMP indicator family, important milestones were the ability to detect rare instances of single action potentials with GCaMP5 (ref. ^26^), and reports of more frequent^23^ albeit still not reliable spike detection^8^ with GCaMP6. The latest iteration of GCaMP optimization, which resulted in several GCaMP8 indicators (variants “f”, “m”, and “s”), was designed to further improve sensitivity and provide faster kinetics compared to its predecessors^27^. Yet, it remains unclear whether calcium imaging data obtained with GCaMP8 should be processed and interpreted differently compared to previous indicator versions. How can methods established for GCaMP6 be adapted for GCaMP8 data? In addition, as GCaMP8 sensors are more linear and exhibit faster rise times compared to previous GCaMP generations^27^, these improved properties raise questions about the consequences for practical applications: For instance, does the smallest detectable calcium transient for GCaMP8 reflect the same underlying spike pattern as for GCaMP6? Furthermore, are there specific advantages of GCaMP8 indicator properties for particular experimental applications?

In this study, we systematically address these questions, taking advantage of existing datasets with simultaneous calcium imaging and electrophysiological juxtacellular recordings of spikes from the same neurons (“ground truth datasets”). We investigate how spike inference based on a deep learning-based method (CASCADE, ref. ^14^) and on model-based approaches (OASIS, MLSpike, refs. ^13,15^) generalizes from previous calcium indicators to GCaMP8. We take the point of view of the experimenter to explore the limitations and potential advantages of using GCaMP8 together with spike inference as a precise readout of neuronal activity. Our analyses cover various recording conditions, both for excitatory and inhibitory neurons, and span a range of recording noise levels that are typical for in vivo population calcium imaging. We find that algorithms for spike inference adapted for GCaMP8 make an important step towards more accurate and interpretable inference of the true spike patterns by taking into account the improved linearity of the “s” and “m” variants of GCaMP8and their ability to pick up single action potentials.

## Results

### Generalization of spike inference algorithms to GCaMP8

We first evaluated how well existing algorithms for spike inference, which had been optimized on GCaMP6 data, were able to generalize to GCaMP8. To enable this evaluation, we used the simultaneously acquired calcium imaging data and electrophysiological juxtacellular recordings from the same cells (“ground truth datasets”) provided with the original GCaMP8 study^27,28^. This dataset (Fig. 1a,b) consists of ground truth recordings obtained from layer 2/3 pyramidal neurons in visual cortex of anesthetized mice during spontaneous neuronal activity or during visual stimulation (36 neurons for GCaMP8f, 42 neurons for GCaMP8m, 39 neurons for GCaMP8s, plus 22 neurons for GCaMP7f; after quality control, see Methods). Compared to previous ground truth recordings from L2/3 pyramidal neurons in mouse cortex^8,23,29^, electrophysiological properties were similar, with a modest increase of the burst propensity (Fig. Supp. 1-1). We complemented this GCaMP8-dataset with an existing compound ground truth dataset using GCaMP6 indicators (Fig. 1b, 125 neurons; described in ref. ^14^). To render evaluations comparable across datasets, we resampled all ground truth datasets to a consistent frame rate (30 Hz unless otherwise indicated) and added Gaussian noise to achieve a specific standardized noise level (see Methods for details). This procedure allowed us to create, from the same ground truth data, low-noise recordings (standardized noise levels ∼2) typical for high-quality population recordings of up to 100-200 neurons^14,30^ and high-noise recordings (noise levels ∼8) typical for recordings with lower SNR or from a large number of neurons (>1000).

**Figure 1.**
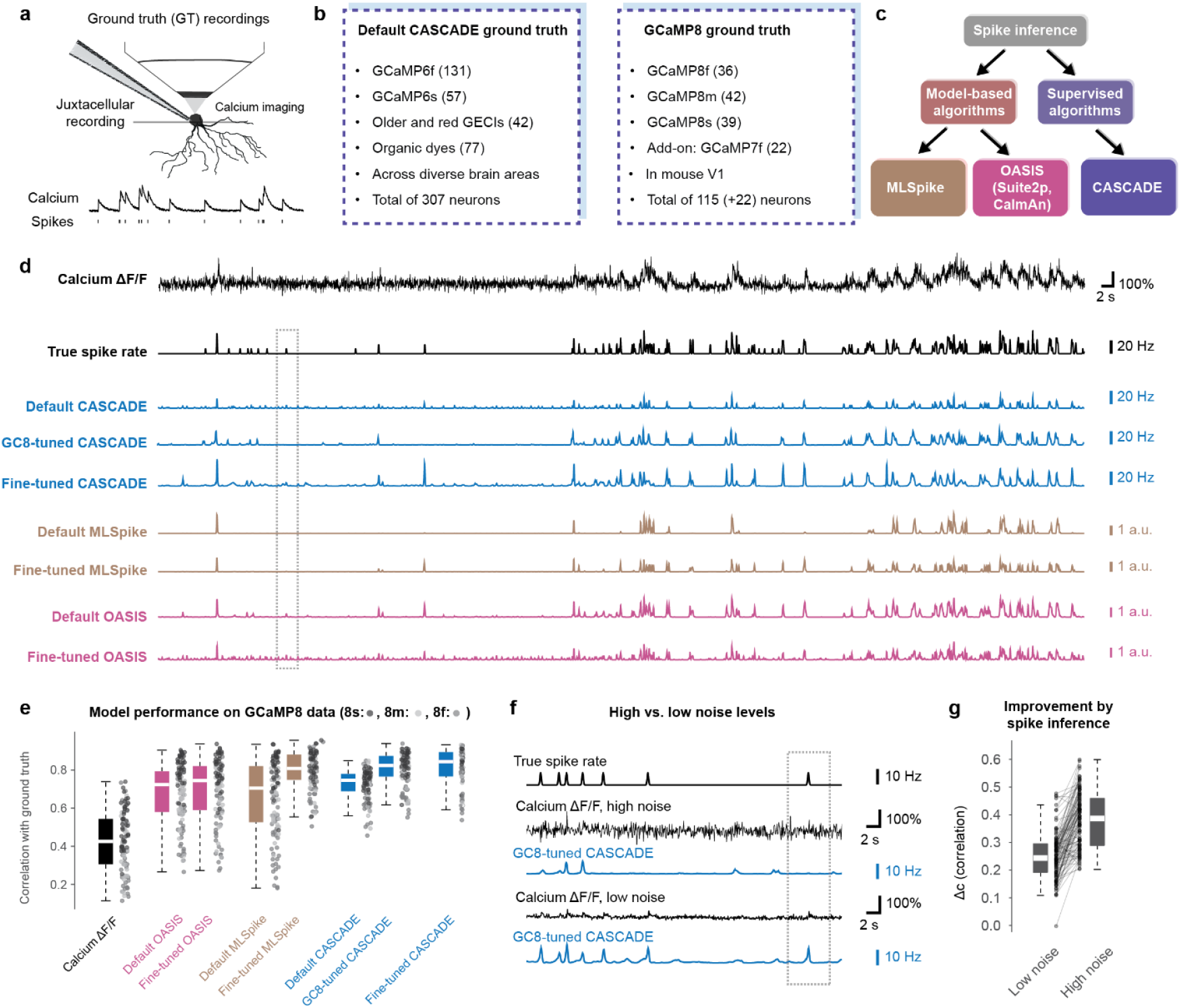
Benchmarking of spike inference algorithms for GCaMP8. **a**, Scheme of ground truth recordings, obtained by simultaneous juxtacellular electrophysiology and two-photon calcium imaging. **b**, Overview of calcium indicator datasets from a database prior to GCaMP8 (left, ref. ^14^) and datasets from the GCaMP8 study (right, ref. ^27^). **c**, Overview of three commonly used algorithms for spike inference. **d**, Example of spike inference from calcium imaging data (top trace, resampled at 30 Hz, at a standardized noise level of “8”, GCaMP8m), with ground truth spike rate (black) as well as spike rates inferred by different variants of the CASCADE algorithm (blue), OASIS (pink) and MLSpike (brown). GC8-tuned models were trained with the ground truth available across all GCaMP8 variants, while fine-tuned models were trained based only on the variant of interest (e.g., GCaMP8m). The dashed box highlights an isolated action potential. Spike inference for the same recording but resampled at lower noise levels is shown in Fig. Suppl. 1-1. **e**, Quantification of model performance across all GCaMP8 ground truth data resampled at 30 Hz and with standardized noise level of “8”. **f**, Example of spike inference from the same ground truth, resampled at high vs. low noise levels. Dashed box as in panel (a). **g**, Improvement of spike inference (increase of correlation, Δc, compared to using ΔF/F as a proxy for neuronal spike rate) for low and high noise levels.

We evaluated several standard algorithms that are used to infer spiking activity from fluorescence traces (Fig. 1c). First, we used the raw ΔF/F traces as a baseline for the benchmark. Second, we applied OASIS, a widely used method implemented in Suite2p^31^ and CaImAn^32^ as the simplest method for spike inference. Third, we used MLSpike^15^, which has consistently been shown to be a state-of-the-art model-based method for spike inference^14,18^. For the model-based methods (OASIS and MLSpike), we provide benchmarking results for the default parameters, but in addition we also analyzed how to adjust model parameters to optimize spike inference for GCaMP8 data (“fine-tuned” models; see Methods). Fourth, we included CASCADE^14^, a supervised method based on deep learning that was trained on a large and diverse set of calcium indicators, but with a focus on GCaMP6. The published version, hence referred to as “Default CASCADE”, has been shown to outperform other existing approaches including MLSpike. Finally, using GCaMP8 ground truth data, we trained new CASCADE models for GCaMP8 data (termed “GC8-tuned CASCADE”) and for each GCaMP8-variant specifically (“Fine-tuned CASCADE”, e.g., “GC8s-tuned CASADE” for GCaMP8s). We then compared each model’s performance in a cross-validated manner on GCaMP8 data using the correlation with ground truth spike rates, obtained by temporal smoothening of the discrete spikes from juxtacellular recordings, as the main performance metric (Methods; Fig. 1d-e).

In general, all models outperformed the baseline set by raw ΔF/F for both high-noise (Fig. 1) and low-noise recordings (Fig. Suppl. 1-2). With default parameters, OASIS and MLSpike performed reasonably well, but their performance was improved by fine-tuning (Fig. 1e). This improvement was modest for OASIS (from 0.72 [0.58; 0.79] to 0.74 [0.59; 0.82], median correlation with inter-quartile ranges [IQR] across 105 neurons) but more prominent for MLSpike, for which more parameters can be optimized (from 0.71 [0.53; 0.82] to 0.81 [0.75; 0.88]). The optimal performance was relatively robust against parameter choices both for OASIS (Fig. Suppl. 1-3) and MLSpike (Fig. Suppl. 1-4). Interestingly, the optimal decay time parameters found for the GCaMP8 variants (0.45-0.70 s for OASIS and 0.3-0.8 s for MLSpike) were much larger than the measured decay time constant of the indicator (0.05-s, ref. ^27^). This discrepancy suggests that the decay constants in these model-based inference methods act as heuristic parameters – values that improve spike inference accuracy – rather than as biophysical parameters that accurately describe the calcium indicator’s true kinetics.

Default CASCADE performed better (0.75 [0.69; 0.78]) than the default version of OASIS and MLSpike (p = 2.3e-4 and 8.6e-4, repeated-measures ANOVA with Tukey-Kramer correction) but was surpassed by the fine-tuned version of MLSpike (p = 6.0e-8). GC8-tuned CASCADE, on the other hand, performed as well as the fine-tuned MLspike (0.82 [0.77; 0.87], p = 0.83) and outperformed all other appoaches (p = 6.0e-8 for all other comparisons). Fine-tuning with each specific GCaMP8 improved the performance compared to fine-tuned MLSpike or GC8-tuned CASCADE only to a minor extent, if at all (0.84 [0.77; 0.89]; p = 0.62 and 0.95) (Fig. 1e). We noticed that CASCADE trained on GCaMP8 data also performed very well for the small ground truth dataset available for GCaMP7f (Fig. Suppl. 1-5). In addition, we noted that spike inference yielded higher correlation with ground truth for GCaMP8 compared to GCaMP6 (Fig. Suppl. 1-6). However, this result requires careful interpretation due to potentially different recording conditions. Interestingly, the retrained models were not universally better; they performed better for GCaMP8 data but worse for datasets with previous indicators (Fig. Suppl. 1-2c). These results demonstrate that spike inference from calcium imaging data with GCaMP8 benefits from model parameter optimization or supervised training performed on specific GCaMP8 data.

The quality of spike inference from two-photon calcium imaging data is limited not only by the indicator and the inference model but also by the recording quality and in particular the shot noise level^33,34^. To assess this dependence, we evaluated all models for both low and high noise levels. We found no major difference in the relative performances of the different algorithms (Fig. 1e, Fig. Suppl. 1-1b). Isolated spikes, which were often missed by CASCADE for the high-noise condition, were more reliably detected for low-noise data (Fig. 1f), as one would expect. Furthermore, spike inference yielded greater improvements in terms of the correlation-based performance for noisier data, reflected by larger performance increases relative to the baseline performance (ΔF/F) (improvement Δc = 0.24 [0.19; 0.3] across neurons for low noise, Δc = 0.39 [0.29; 0.46] for high noise) (Fig. 1f,g). This finding expands on the well-established observation that spike inference effectively denoises calcium imaging data^13,14^

In conclusion, spike inference generalized well from previous indicators to GCaMP8 but fine-tuning these methods for GCaMP8 further enhanced the performance. To support the wider adoption of these optimized algorithms, we provide the best-fit parameters for each GCaMP8 variant in the Methods section for OASIS and MLSpike, and pretrained models for CASCADE online (https://github.com/HelmchenLabSoftware/Cascade).

### The effect of nonlinearities on high-frequency spike patterns

It has been previously shown that GCaMP8 variants, in particular GCaMP8m/s, are more linear calcium sensors, especially when compared to GCaMP6 variants^27^. We performed an in-depth quantitative analysis of this relationship for existing ground truth datasets, using transfer functions between true and inferred spike rates (Supplementary Note 1). We observed that the nonlinearity of GCaMP6 sensors is captured by Default CASCADE, resulting, however, in distorted predictions when applied to GCaMP8 data. In contrast, CASCADE models trained with GCaMP8 data exhibited a higher linearity, demonstrating that the calcium indicators GCaMP8s and GCaMP8m, as well as the models trained with these data, behave much more linearly than previous indicators and models (more details in Supplementary Note 1).

Next, we wanted to understand how specific spike patterns and not only transfer functions are affected by spike inference specifically for GCaMP8. The sigmoidal relationship of indicator fluorescence and intracellular calcium concentration is often described by the Hill coefficient *n*, which determines the sigmoidal shape, and the dissociation constant *K*_*d*_, which determines the inflection point of the sigmoid in relation to the baseline calcium concentration (ref. ^9^). For GCaMP6 indicators, with *K*_*d*_ higher than the baseline calcium concentration of the neuron, this nonlinearity typically results in larger fluorescence increases when more calcium is already bound by the indicator (Fig. 2a). Based on this more nonlinear behavior, we reasoned that spikes which occur later during a high-frequency event will result in a prominent and disproportionately large fluorescence increment for GCaMP6. We hypothesized that this behavior will not occur for GCaMP8 indicators, which are supposed to behave more linearly around the neuronal baseline calcium level due to their lower *K*_*d*_ values. To provide an intuition of this history-dependent fluorescence change for GCaMP6, we demonstrate that this effect can be obtained with a typical sigmoidal model^6^ of GCaMP nonlinearities (Fig. 2b-d; see Methods). Another mechanism that potentially contributes to such a history-dependent nonlinear effect of a calcium indicator is the recently demonstrated use-dependent slowing of calcium indicators^35^, although the role of this effect has not yet been investigated for GCaMP6s and GCaMP8m/s.

**Figure 2.**
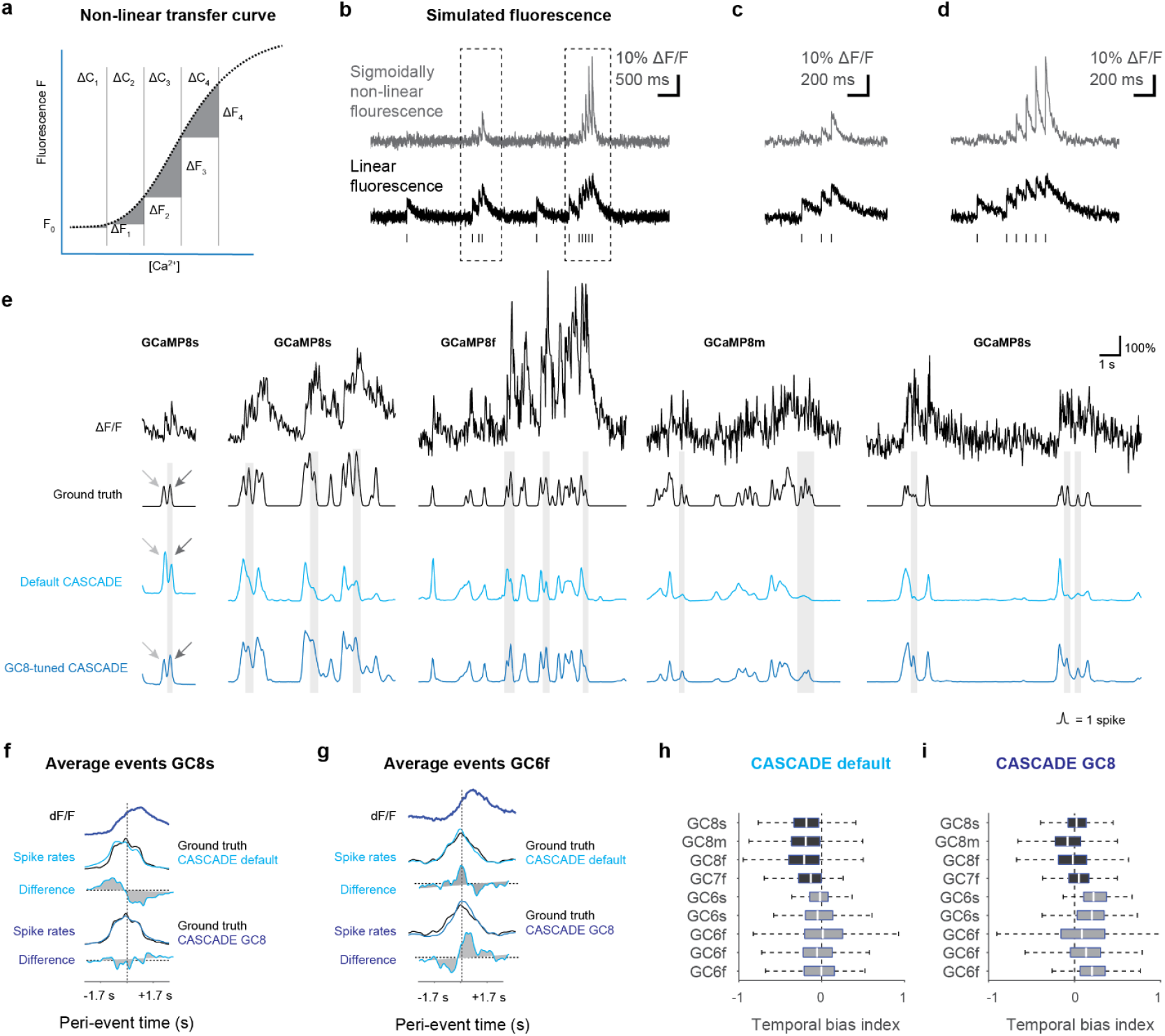
Improved spike inference during high-frequency spike events for fine-tuned CASCADE models. **a**, Scheme: fluorescence change ΔF as a function of calcium concentration change ΔC for a nonlinear indicator such as GCaMP6. For such a nonlinear relationship, the first action potential (ΔC_1_) might go unnoticed (ΔF_1_), while subsequent action potentials will trigger a much larger response (ΔF_2_ or ΔF_3_). **b**, Illustration of the effects of a sigmoidal nonlinearity as shown in (a) on high-frequency events using simulated data. ΔF/F amplitudes for later spikes are exaggerated compared to early spikes for a nonlinear indicator (gray) but not for a linear one (black). This property illustrates the nonlinear behavior that is prominent for GCaMP6. **c-d**, Zoom-ins of panel (b), highlighting sub-linear suppression of initial and supralinear amplification of later action potentials for the nonlinear model. **e**, Examples of high-frequency events that show a biased estimation of spike rates inferred from GCaMP8 signals by Default CASCADE but not GC8-tuned CASCADE. The earlier time points of a high-frequency event are exaggerated by Default CASCADE (light gray arrow) while later time points of a high-frequency event are suppressed (dark gray arrow). The gray shading highlights cases where Default CASCADE underestimates the spike rate for trailing edges of high-frequency events. **f**, From top to bottom: average high-frequency event traces for ΔF/F; spike rates obtained with Default CASCADE and the difference compared to true spike rates; spike rates obtained with GC8-tuned CASCADE and the difference compared to true spike rates. The difference plot illustrates the bias induced by Default CASCADE but not GC8-tuned CASCADE. Data from the GCaMP8s ground truth dataset. Ground truth and inferred spike rates are scaled for best mutual linear fit, and scale bars are therefore omitted. **g**, Same as in (f) but for the GCaMP6f dataset. **h**, Temporal bias as shown in (f) and (g) quantified for each dataset when applying Default CASCADE. Temporal bias index as defined in the main text. A box plot centered around zero indicates low or no bias. **i**, Same as in (h) but when applying GC8-tuned CASCADE.

To analyze the relevance of these effects for spike inference, we focused on CASCADE models, which exhibited the best performance for GCaMP6 (ref. ^14^; Default CASCADE) and GCaMP8 (Fig. 1; GC8-tuned CASCADE). This focus allows for precise comparison between the GCaMP6 and GCaMP8 indicator families, as well as for comparisons between models optimized for these indicators. We visually screened inferred spike rates together with ground truth and noticed systematic distortions of inferred spike rates during high-frequency events, i.e., when multiple spikes and bursts occurred within a short time window. For illustration, we showcase such examples across all GCaMP8 variants (Fig. 2e). For these high-frequency events, Default CASCADE tended to overestimate the spike rate for the initial spikes while underestimating the spike rate for subsequent spike patterns. These systematic mistakes were largely corrected by using GC8-tuned CASCADE (Fig. 2a).

To systematically confirm our initial observation, we detected high-frequency events based on a simple threshold (instantaneous smoothed ground truth spike rate >6 Hz). Then, we extracted the raw ΔF/F trace and inferred spike rates in a time window centered around the maximum true spike rate (“peak time”). We performed this analysis both for GCaMP8 and GCaMP6 data from our joint ground truth database (Fig. 1b). To quantify potential biases towards the initial vs. late phase of such high-frequency events, we computed the difference between normalized true and inferred spike rate. Using the GCaMP8s dataset as an example, the application of Default CASCADE resulted in an overestimation of the spike rate during the initial phase and an underestimation during the later phase of the high-frequency events (Fig. 2b, based on 1218 events across 37 neurons). These biases were eliminated when we applied GC8-tuned CASCADE (Fig. 2f). In contrast, for a GCaMP6f dataset, Default CASCADE resulted in an unbiased estimation of spike rates during high-frequency events, whereas GC8-tuned CASCADE led to an overestimation of late phase-spike rates (Fig. 2g, based on 329 events across 23 neurons).

To summarize this finding across datasets, we computed for each difference curve as in Fig. 2b,c the area under the curve before (*AUC*_*pre*_) and after (*AUC*_*post*_) the peak time and calculated the normalized difference as a temporal bias index (TBI): *TBI* = (*AUC*_*post*_ − *AUC*_*pre*_)/(*AUC*_*post*_ + *AUC*_*pre*_). Default CASCADE resulted in a negative bias for all GCaMP8 datasets (TBI = -0.20 ± 0.02, mean ± s.d. across the 3 datasets) and no visible bias for GCaMP6 data (-0.02 ± 0.03 across the 5 datasets; Fig. 2h). Conversely, GC8-tuned CASCADE exhibited no visible bias for GCaMP8 (-0.03 ± 0.05) but a positive bias for GCaMP6 datasets (0.17 ± 0.05; Fig. 2i). Interestingly, the calcium indicator GCaMP7f positioned itself somewhat between GCaMP6 and GCaMP8 (Fig. 2h-i).

As a summary, our analyses demonstrate the effect of a nonlinear calcium-to-fluorescence relationship on the fluorescence recorded during high-frequency events, as well as on the analysis of such events with spike inference. We find that Default CASCADE effectively compensates for the history-dependent distortions by GCaMP6, resulting in unbiased spike rates around high-frequency events for GCaMP6 data (Fig. 2h), while introducing a systematic bias for GCaMP8 data. In contrast, spike inference optimized for data acquired with the more linear GCaMP8 indicators processed high-frequency events from GCaMP8 data without introducing a bias. These results demonstrate that the distinct nonlinear properties of GCaMP6 and GCaMP8 are reflected in models trained on these ground truth datasets and systematically affect the analysis of prolonged and complex spiking events. Our analysis underscores the need for a GCaMP8-specific optimization of spike inference to avoid history-dependent biases in estimating neuronal activity.

### Single-action potential detection with GCaMP8

A key question in neuronal calcium imaging is whether specific calcium indicators can reveal single action potentials (APs). This question is crucial as the detection of single APs serves as an intuitive benchmark that complements other metrics such as correlation with ground truth. Furthermore, whether the electrical signals underlying fluorescence changes are bursts or isolated APs is highly important for the interpretation of neuronal activity^36–38^. Resolving this issue would make neuronal calcium imaging more interpretable and help to tackle fundamental questions about the nature of the neuronal code.

While it has often been claimed that single APs *can* be resolved using specific algorithms or indicators^10,23,24^, the conditions necessary for single AP detection could often not be achieved in typical experimental conditions^39–41^. Moreover, detectable AP-triggered calcium transients in previous calcium indicators were frequently observed only in a small and not necessarily representative subset of neurons^14,42,43^. We therefore addressed the question in a systematic and comparative approach: can single APs be reliably identified with GCaMP8?

Detection of single APs requires a trade-off between false positive vs. false negative detections. For cortical recordings of pyramidal cell activity, neuronal silence is more prevalent than activity, a condition that can lead to frequent false positives if the detection algorithm is not appropriately balanced. Previous studies used a manually chosen and therefore to some extent arbitrary threshold of false-positive detection rates to evaluate single-AP detection^39,41^. Here, we used CASCADE to automatically choose the best trade-off. CASCADE is trained in a supervised manner to minimize the mean squared deviation from ground truth for the entire recordings. Thus, it takes into account both silent and active periods and the frequency with which these periods occur. The single-AP detection ability of such an algorithm – which is not specifically optimized for single-AP detection – is therefore a benchmark that provides realistic performance metrics.

To assess the detection of single APs, we identified isolated APs and the surrounding calcium traces in ground truth recordings (see Methods). Then, we added a variable amount of Gaussian noise to the calcium traces and inferred spike rates from these traces using a CASCADE model trained for this entire dataset except the tested neuron (*e*.*g*., GC8s-tuned CASCADE for the GCaMP8s data). Strikingly, we found that single APs were almost never detected with GCaMP6f and rarely with GCaMP6s, GCaMP7f or GCaMP8f (Fig. 3a; Fig. Suppl. 3-1). In contrast, GCaMP8m and GCaMP8s demonstrated a relatively good ability to detect isolated APs, which deteriorated, however, with increasing noise levels (Fig. 3a; Fig. Suppl. 3-1).

**Figure 3.**
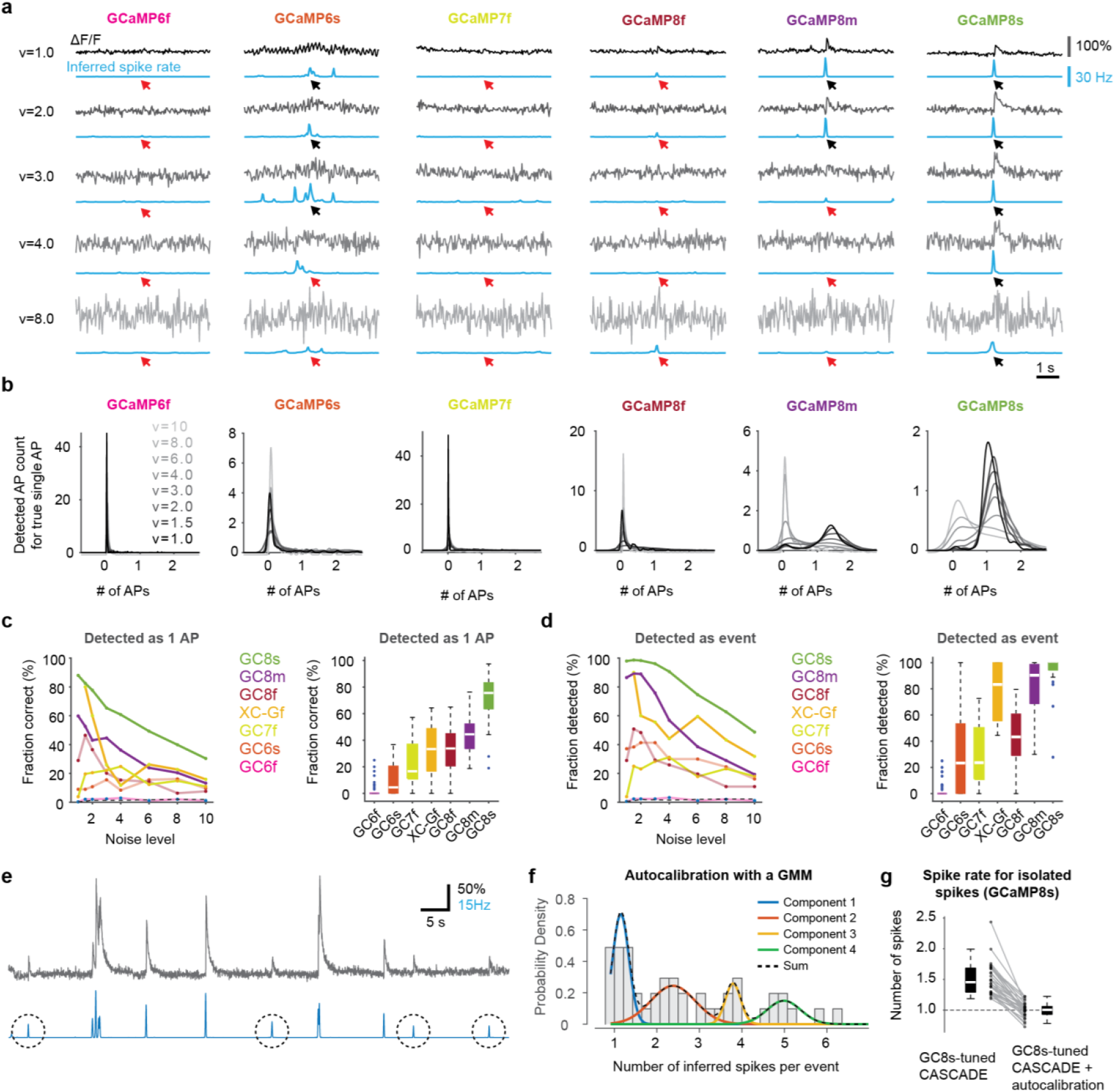
Detection of isolated action potentials from GCaMP8 with spike inference. **a**, Examples of isolated action potentials with the corresponding ΔF/F trace (gray) and the inferred spike rate (blue) for an increasing standardized noise level ν. A low-noise population recording corresponds to ν ≈ 2, a higher-noise population recording to ν ≈ 8. Arrows indicate the time point when the true isolated action potential occurred (red arrow if not detected). Single action potentials are accurately inferred for low noise levels for GCaMP8m/s. **b**, Number of detected action potentials (APs) for a true isolated single AP, plotted as a distribution (kernel density estimate). Shades of gray indicate the different noise levels (lowest noise level, dark grey; highest noise level, light grey). Ideally, the distribution would be narrow and centered around “1” true action potential. **c**, *Left:* Fraction of APs correctly detected as a single AP (spike count >0.5 and <1.5) across datasets and noise levels. *Right*: Fraction of APs per neuron correctly detected as a single AP, averaged across noise levels ν = 2-4 for robustness. **d**, *Left:* Fraction of APs correctly distinguished from noise (spike count >0.5) across datasets and noise levels. *Right:* Fraction of APs per neuron distinguished as event, averaged across noise levels ν = 2-4. **e**, Example of a calcium imaging recording (GCaMP8s ground truth) together with spike rates inferred using GC8s-tuned CASCADE. Isolated minimal events are visually detectable (circles). **f**, Histogram of inferred spike numbers for isolated events prior to auto-calibration, together with a Gaussian mixture model (GMM) fit to the underlying data. The unitary amplitude, defined as the mean of the first Gaussian component (blue), is used for auto-calibration. **g**, Auto-calibration with unitary amplitudes derived from GMM fits decreases the overestimation of the spike rate for isolated spike events and brings back outliers. (But see also Fig. Suppl. 1-3.)

To quantify these observations, we evaluated the distribution of inferred spike rates for each single-AP event using a kernel density estimate. We found that this distribution was centered around zero for GCaMP6f, GCaMP6s, GCaMP7f and GCaMP8f. This finding was even more pronounced for high noise levels, reflecting the inability to detect single APs (Fig. 3b). We note, however, that some degree of variability exists across datasets obtained with the same indicator (Fig. Suppl. 3-2). Thus, for these indicators, AP-evoked calcium transients are in most cases indistinguishable from noise under typical conditions. Consequently, the supervised algorithm decides to make the conservative estimate of “0” spikes. Interestingly, this situation is different for GCaMP8m and GCaMP8s, for which the distribution of inferred spikes for isolated single true APs was centered around “1” spike for low or moderate noise levels (Fig. 3b). For GCaMP8m, the distribution shifted back to the “0” bin for intermediate noise levels, whereas this happened for GCaMP8s only at the highest noise levels. We confirmed that these results did not depend on the use of CASCADE for analysis and could also be observed when using MLSpike (Fig. 3-3).

To summarize these results across different noise levels, we quantified the fraction of true isolated APs that can be detected as a single AP (inferred spike rate between 0.5 and 1.5 APs). In addition, we also quantified the fraction of such events that contained one true AP and were detected at all as a spiking event (inferred spike rate >0.5 APs). As expected from the distributions shown in Fig. 3b, GCaMP8s emerged as the best available indicator, with a fraction of correct detections of >80% and a fraction of detected events close to 100% for the lowest noise levels (Fig. 3c,d). GCaMP8m also exhibited high detection rates that were, however, clearly lower compared to GCaMP8s, followed with some distance by GCaMP8f, GCaMP7f, GCaMP6s and – with a detection fraction of close to 0% across all noise conditions – by GCaMP6f. The calcium indicator XCaMP-Gf deserves a special mention: although the available dataset is relatively small, in particular for low noise levels (see Suppl. Table 1), single APs could be detected more reliably compared to GCaMP6 variants and GCaMP7f, probably reflecting the more linear behavior of the XCaMP indicator family^25^.

For the analyses in Fig. 3b and 3c,d-left, AP events were pooled across all neurons. To exclude a potential bias introduced by many events from a small subset of neurons, we additionally computed detection rates averaged across events for each neuron (Fig. 3c,d-right). These quantifications confirmed the excellent performance of GCaMP8s for the detection of isolated APs and the relatively good performance of GCaMP8m and XCaMP-Gf. This analysis further corroborated the finding that GCaMP6 data are largely incapable of detecting isolated APs in these cortical recordings.

Next, we analyzed how the ability to detect single spikes may affect the overall performance of spike inference as quantified in Fig. 1. We reasoned that, if a calcium indicator cannot reliably detect isolated APs, spike inference must rely on high-frequency events and bursts and will therefore perform worse for less bursty neurons. Indeed, we found that the coefficient of variation of inter-spike intervals of a neuron (Fig. Suppl. 1-1) exhibited a significant correlation with the performance of spike inference for GCaMP8f (correlation c = 0.42, p = 0.01, n = 36 neurons; performance for fine-tuned CASCADE) and GCaMP6 (c = 0.43, p < 10^-5^, n = 98 neurons from refs. ^8,23^), but not for GCaMP8m (c = 0.22, p = 0.17, n = 42) and GCaMP8s (c = 0.24, p = 0.14, n = 39), indicating that spike inference from cortical principal cells relies on bursts of APs to a larger extent for GCaMP6 and GCaMP8f than for GCaMP8m/s. Together, these results provide guidance for the interpretation of calcium signals as bursts vs. single APs from these diverse indicators.

Next, we wondered whether the reliable detection of isolated APs could enable more precise spike inference. By using a quantal analysis to identify the smallest detectable calcium transient corresponding to a single AP, one could potentially rescale the transfer function of each individual neuron with a constant factor to match the identity line (cf. Supplementary Note 1). Previous attempts of such an “auto-calibration” by quantal analysis do exist^15,43^ but have not been successfully applied^18^, also due to the fact that isolated APs are not reflected in reliably detectable calcium transients for GCaMP6 (Fig. 3b-d). With the improved ability of GCaMP8m/s to detect isolated APs, we performed an auto-calibration upon detection of discrete isolated events in a proof-of-concept analysis using the GCaMP8s dataset (Fig. 3e). To identify discrete events, we created a histogram of inferred spike rates and used Gaussian mixture modeling (GMM) to extract the unitary response (Fig. 3f, see Methods for details). Crucially, the unitary response enabled us to estimate single APs more accurately and in a less biased manner by linear rescaling of the inferred spike rates (Fig. 3g, Fig. 3-4). We therefore show as proof of concept that auto-calibration based on single AP-evoked calcium transients is possible. However, we observed that such a simple linear re-scaling degraded the performance for high firing rate regimes (Fig. 3-4b,d), most likely due to the residual nonlinearity of the GCaMP8 indicator. Using subtractive re-scaling as a less principled approach worked better but resulted in higher dispersion of inferred spike rates for low true spike rates (Fig. 3-4c,d). This analysis demonstrates the detrimental effect of the still existing nonlinearities of GCaMP8 and calls for more advanced algorithms for auto-calibration that infer not only unitary response magnitudes but also nonlinearities of calcium indicators from a calcium recording without ground truth.

### Indicator linearity enables single-action potential detection

GCaMP8 was introduced as particularly advantageous due to its fast kinetics and large overall ΔF/F responses. However, based on our analyses (Figs. 2 and 3), we hypothesized that the ability to detect single action potentials – one of most advantageous properties of GCaMP8m/s – comes primarily from the more linear behavior of these indicators when isolated or few APs occur. To shape an intuition for this idea, we used existing ground-truth datasets of different indicators to extract each neuron’s linear response function to the average AP (Methods). Then, we used this linear response to generate ΔF/F traces based on measured spikes, adding Gaussian noise to match experimental conditions, simulating linear indicator behavior (“linearized ground truth dataset”). We use this approach to investigate the difference between linear and nonlinear indicator behavior while leaving kinetics, overall response magnitude and noise level unaffected. The comparison of measured vs. simulated ΔF/F highlights the most striking consequences of the nonlinearity of GCaMP6 (Fig. 4a): (1) bursts appeared exaggerated for nonlinear but not for linear indicator behavior, (2) single APs were undetectable with nonlinear but detectable with linear behavior under otherwise identical conditions, and (3) the onset of high-frequency events as described in Fig. 2 was misrepresented under nonlinear but not linear indicator behavior. Importantly, none of these differences could be observed for the more linear indicator GCaMP8s (Fig. 4b). These comparisons demonstrate that indicators like GCaMP6f/s, with unchanged indicator kinetics and response amplitudes but with linear behavior, would be able to resolve single APs.

**Figure 4.**
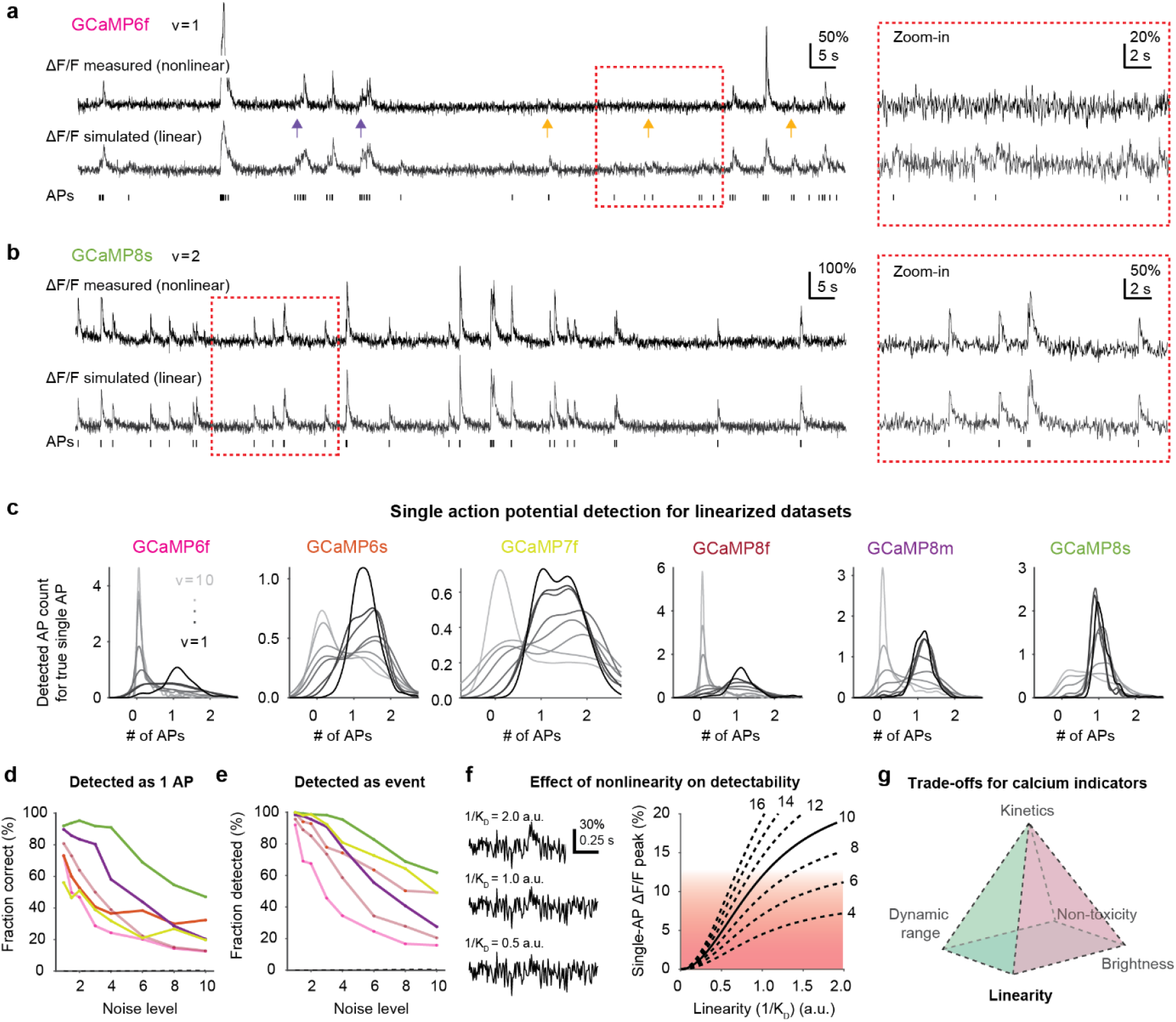
Indicator linearity enables the detection of isolated action potentials. **a**, Measured (top) and simulated linearized (bottom) ΔF/F trace for the same recording with GCaMP6f. True action potentials (APs) are shown as ticks below. The simulation is based on the linear kernel, resulting in ΔF/F traces with, on average, the same response magnitude as the measured data and with the same noise level (standardized noise level ν = 1), but with a linear behavior of ΔF/F vs. spiking. Orange arrows highlight instances of isolated APs with visible deflections in the simulated linearized but not in the measured nonlinear case. Blue arrows highlight instances of history-dependent effects for linear vs. nonlinear indicators as described in Fig. 2. **b**, Same as in (a) but for a recording with GCaMP8s. No striking difference between simulated and measured ΔF/F traces can be observed. **c**, Number of detected APs for a true isolated single AP, as in Fig. 3b. Notably, for low noise levels (dark colors), single APs can be detected also for GCaMP6f/s despite unchanged noise levels and signal amplitudes compared to experimental data (cf. Fig. 3b). **d**, Fraction of APs correctly detected as a single AP across datasets and noise levels for linearized simulated data. **e**, Fraction of APs correctly distinguished from noise across datasets and noise levels for linearized simulated data. **f**, Data simulated with purely exponential kernels, illustrating the effect of linearity on the detectability of isolated APs. The nonlinearity is controlled with the K_D_ value of a Hill-type nonlinearity with exponent n = 2. *Left:* Example transients for a single AP with identical noise levels and baseline fluorescence responses but changed nonlinearity. *Right:* Peak amplitude of single AP-evoked ΔF/F transients as a function of 1/K_D_ (arbitrary units) as a parameter to control simulated indicator linearity. Only conditions that surpass the ΔF/F noise of ∼10% (red shading), typical for standardized noise levels of ν = 2, will result in detectable ΔF/F transients for single APs. Dashed lines indicate simulations with different levels of baseline fluorescence responses (indicator brightness x dynamic range, in arbitrary units) prior to the nonlinearity. Even for the highest baseline fluorescence responses, single APs cannot be detected if the indicator is highly nonlinear (left quadrant of the graph). **g**, Illustration of trade-offs for calcium indicator design, highlighting the importance of indicator linearity, also to enable detection of single APs.

To quantify this initial observation, we repeated the analysis described in Fig. 3b-d, but with simulated “linearized” data (Fig. 4c-e). We observed that all linearized indicators were able to detect single action potentials with considerable reliability at low noise level, and performance degraded gradually as a function of the response amplitude of each indicator. Importantly, the single-AP detection power of “linearized” GCaMP8s was only slightly higher than for the experimentally measured GCaMP8s dataset, indicating that the indicator itself is already relatively linear for the low firing-rate regime.

To illustrate the isolated effect of linearity more systematically, we simulated a spike-to-fluorescence relationship with a standard Hill nonlinearity and varied the *K*_*D*_ value that controls the inflection point of the transfer function, which is one possible method to artificially control the nonlinearity (Methods). This analysis demonstrates that the linearity of an calcium indicator can affect the ability to detect single APs in a nonlinear manner (Fig. 4f). It also demonstrates that even for bright sensors with large dynamic range, nonlinear behavior can result in non-detectable single APs, further underlining the necessity of linear calcium indicators.

Together, these findings demonstrate that the linearity of calcium indicators, especially in the low-firing regime, is key to enabling the detection of single action potentials. Linearity as an optimization metric should therefore be an essential criterion both for researchers who select calcium indicators for their experiments and for engineers who design the next generation of calcium sensors (Fig. 4g).

### Online spike inference with fast rise times

A prominent feature of GCaMP8 is its faster fluorescence rise time compared to previous genetically encoded calcium indicators^27^. What are the implications of this effect for spike inference from calcium imaging data? To study this question, we first used linear deconvolution (Methods) to extract the fluorescence kernel evoked by the average AP (Fig. 5a). This linear kernel reflects the average fluorescence transient triggered by any AP, from both complex and isolated events (which is not equal to the fluorescence transient triggered by isolated APs; see Methods). From this extracted kernel, we determined from the original recordings the half-maximum rise time for each neuron across all datasets.

**Figure 5.**
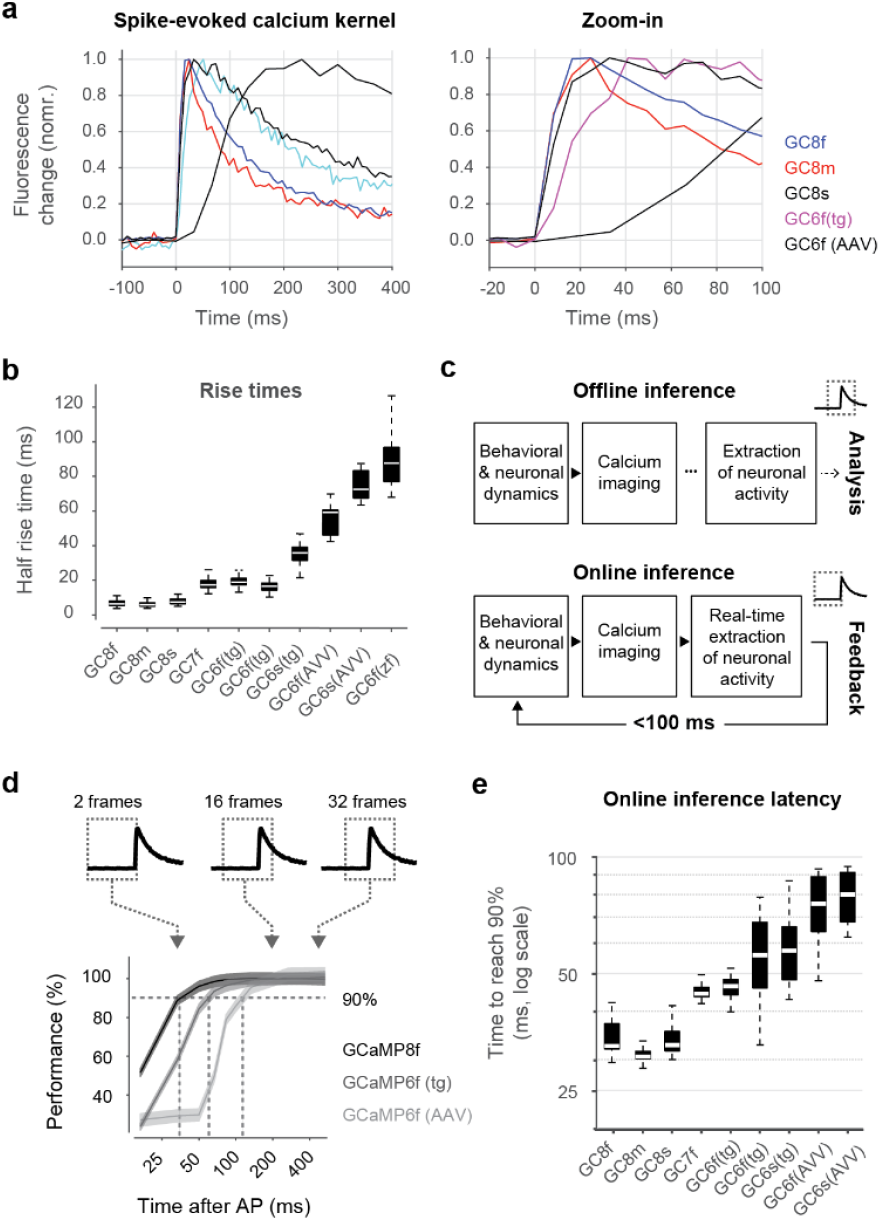
Improved real-time spike inference for GCaMP8 due to faster onset rise times. **a**, Spike-evoked calcium transients for a selected subset of datasets to illustrate differences of overall kinetics (left) and of rise times (zoom-in, right). **b**, Rise times to half of maximum across indicators. Box plots represent distributions across neurons for each dataset. As in panel (a), analyses are performed at the original imaging rate (122 Hz for GC8 and GC7, 158 Hz for GC6 (transgenic), 60 Hz for GC6 (AAV) and 30 Hz for GC6 (zf, zebrafish). **c**, Schematic depiction of offline processing with offline inference (top) and online processing with online spike inference (bottom). In the latter case, processing should be completed within approx. 100 ms, taking into account only calcium signals recorded to this point in time. **d**, Illustration of online spike inference, with a limited number of imaging frames available from the time point after the occurrence of a spike. The resulting performance (% of maximal performance for this dataset) increases sigmoidally as a function of time after the action potential used for inference. For each condition (2 frames, 5 frames, 16 frames, etc.), a new CASCADE model is trained to optimally take advantage of the limited amount of information. Sigmoidal performance dependencies are shown for three example datasets. Analyses are performed on calcium imaging data resampled at 60 Hz. **e**, Time to reach 90% of the maximal performance, distribution across neurons shown for each dataset, for low noise (standardized noise level 2). Comparison across noise levels in Figure 5-1.

As expected, we observed fast rise times for GCaMP8 variants (median rise time with inter-quartiles: 6.4 ms [5.3; 8.0] for GCaMP8f, 5.8 ms [5.0; 7.1] for GCaMP8m, 7.4 ms [6.3; 9.2] for GCaMP8s) and slightly slower rise times for GCaMP7f (17 ms [16; 20]) (Fig. 5b). The results for GCaMP6, however, need to be unpacked more carefully. In agreement with previous analyses^27^, the AAV-mediated GCaMP6 variants exhibited relatively slow rise times (59 ms [46; 61] for GCaMP6f and 72 ms [67; 83] for GCaMP6s). However, datasets based on transgenic expression of GCaMP6f and, to a lesser extent also GCaMP6s, resulted in surprisingly fast rise times (19 ms [17; 21] and 17 ms [14; 18] for GCaMP6f datasets, 36 ms [31; 39] for GCaMP6s). It seems possible that this observation can be explained by the lower expression levels for transgenic as opposed to AAV-based strategies, although the mechanism behind this effect is not clear. Interestingly, a dataset based on transgenic expression of GCaMP6f in zebrafish (from ref. ^14^) resulted in relatively slow rise times (88 ms [77; 97]), likely due to different temperature conditions (room temperature for zebrafish vs. body temperature for mice). These analyses show that rise times are significantly faster for GCaMP8 but may also be affected by other factors like expression strategies, calcium buffering and temperature.

We reasoned that the fast rise times of GCaMP8 could be particularly advantageous for low-latency closed-loop processing. In such experiments, calcium signals are recorded and processed in real-time, enabling the use of feedback signals based on the activity of a defined set of neurons (Fig. 5c). Such paradigms have been applied to study learning and neuronal plasticity^44,45^ and are also key for the development of brain-machine-interfaces^46^. A primary limitation of these feedback loops is the latency from the action potential to the feedback signal derived from this very action potential. This latency includes the action potential-induced calcium influx, binding to the fluorescent calcium sensor, fluorescence imaging with a microscope, extraction of the fluorescence trace from the recorded data, and the time for an algorithm to detect spiking from the extracted trace. To be useful for brain-machine-interfaces and to study learning, such a feedback loop – with all processing steps including motion correction, ROI extraction and spike detection – should have a latency of <100 ms^46^. We hypothesized that the fast rise time of GCaMP8 might enable detection of spike events matching this requirement.

To systematically analyze the impact of calcium indicator rise time on online spike inference, we trained for each dataset (Fig. 1b) multiple CASCADE models that had access to a variable amount of time points after the current time point of interest, which we term “integration time” (see Methods; Fig. 5d). These models were trained and evaluated at a relatively high sampling rate of 60 Hz to enable comparisons at high temporal resolution. We quantified the performance of these models (correlation with ground truth), which resulted in a sigmoidal curve as a function of integration time, with performance increasing for prolonged integration times (Fig. 5d). We assessed the integration time necessary to achieve 90% of the optimal performance (Fig. 5d,e). This integration time indicates how long one needs to wait after a spiking event to achieve this performance criterion. For low-noise calcium imaging data (standardized noise level of “2”), GCaMP8 data exhibited the lowest 90% integration times (32 ms [32; 37] for GCaMP8f, 31 ms [30; 32] for GCaMP8m, 33 ms [32; 35] for GCaMP8s; median [inter-quartile range] across neurons), closely followed by GCaMP7f (44 ms [44; 46]) and transgenically expressed GCaMP6 (46 ms [44; 48] and 56 ms [46; 68] for the GCaMP6f datasets, 57 [48; 66] for GCaMP6s), and, finally, by AAV-induced GCaMP6 (76 ms [64; 89] for GCaMP6f, 80 ms [68; 91] for GCaMP6s). These 90% integration times reflect the ranking of indicator rise times (Fig. 5e, left panel; cf. Fig. 5b). Interestingly, no clear advantage of GCaMP8f compared to the slower variants of GCaMP8 could be observed. For higher noise levels of calcium recordings, the required integration times increased and diminished the advantage of GCaMP8 compared to GCaMP7f and the transgenic variant of GCaMP6f (Fig. S5-1). For example, at a standardized noise level of “8”, which is more typical of large-field-of-view recordings with fewer pixels per neuron, integration times around 50 ms were necessary for the GCaMP8 variants as well as for GCaMP7f. In summary, our findings demonstrate that online spike inference will benefit directly from the fast rise times of GCaMP8. A similar, albeit less pronounced, positive effect may also be achieved by calcium indicators like GCaMP6f that are induced via transgenic expression instead of AAVs. However, only for GCaMP8 indicators and for low noise levels, the integration times were below the desired latency of 100 ms with a considerable margin of safety.

### Spike inference for interneurons

Calcium indicators and algorithms for spike inference are typically tested with pyramidal cell data from mouse cortex, and it is unclear how spike inference will generalize to other brain regions or different cell types. A particularly striking example are cortical fast-spiking interneurons, which have been repeatedly shown to exhibit firing patterns and calcium responses that are distinct from excitatory pyramidal cells^14,47^. Here, we address this issue by taking advantage of ground truth recordings obtained from cortical fast-spiking interneurons identified by electrophysiological spike waveform and spike rate (total of 17 neuronal ground truth recordings; from ref. ^27^).

First, we characterized the interneuron ground truth and found that average spike rates were widely distributed across neurons but on average much higher than for excitatory ground truth recordings (p = 6×10^-6^, Wilcoxon rank-sum test; median spike rate with inter-quartiles 4.4 Hz [2.7; 10.5] for interneurons, 0.72 Hz [0.39; 1.4] for excitatory neurons; Fig. 6a). Calcium transients evoked by the average spike displayed lower amplitudes in interneurons compared to excitatory neurons by an order of magnitude (Fig. 6b,c), in line with previous studies on other calcium indicators^48,49^ and on GCaMP8 (ref. ^27^). The decay appeared slightly slower for interneurons, possibly due to the recently discovered use-dependent slowing of calcium binding^35^. Furthermore, interneurons showed a significantly reduced bursting propensity compared to cortical pyramidal cells as quantified by the coefficient of variation of inter-spike intervals (p = 0.008, Wilcoxon rank-sum test; median coefficient of variation with inter-quartile intervals: 2.0 [1.5; 3.3] for interneurons and 3.1 [2.4; 3.9] for pyramidal cells; Fig. 6d). In a recent study, we have shown that a reduced bursting propensity in non-cortical neurons (dorsal horn of spinal cord) led to diminished performance of spike inference algorithms designed for cortical excitatory neurons^42^. As for these spinal cord neurons, calcium signals recorded from fast-spiking interneurons often change only slowly, with gradual changes reflecting changes in continuous spiking rather than sudden bursts of spikes. We noticed that “default” model-based algorithms, designed for cortical excitatory neurons, often captured only the large and prominent transients that reflect burst activity (Fig. 6e). The performance of default OASIS and MLSpike was only slightly better than using raw ΔF/F. To optimize both algorithms for calcium imaging with interneurons (“fine-tuned” algorithms), we performed grid search across parameter space and observed a significant performance increase in both instances. However, the results of this optimization process also suggest that the algorithms are not well-suited to deal with interneurons: Fine-tuning of OASIS led to a decay time parameter close to zero (see Methods), resulting in an inferred spike rate that closely resembled the original ΔF/F trace (Fig. 6f). Fine-tuning of MLSpike, on the other hand, led to a very high decay time parameter (see Methods), resulting in a stronger emphasis on spike rate changes (Fig. 6e). These optimizations show how spike inference can go awry and how parameter values originally related to physiological processes can become arbitrary when the algorithms are optimized for data that behave very differently from the data used to design the algorithms in the first place.

**Figure 6.**
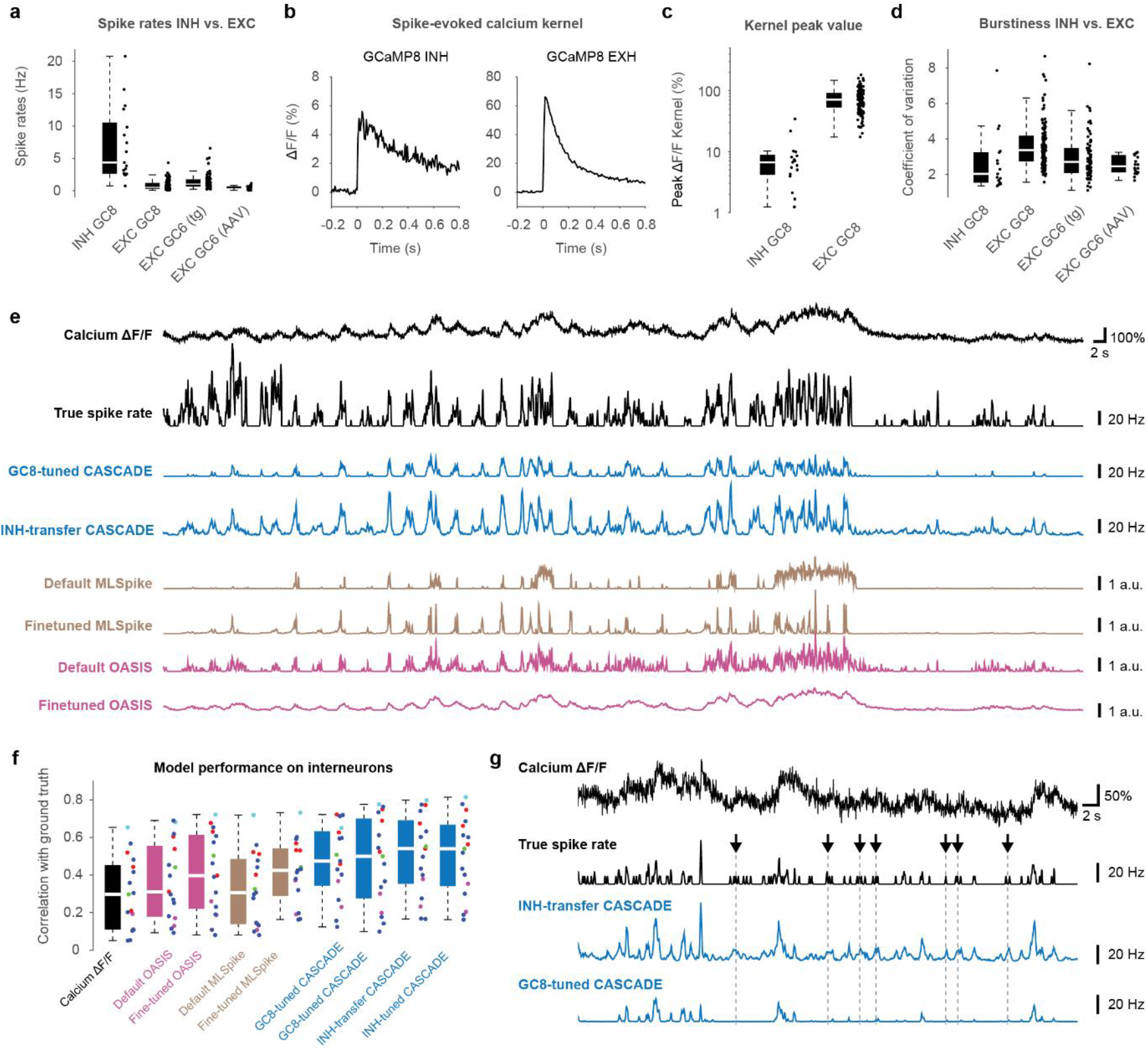
Benchmarking of spike inference algorithms for inhibitory neurons. **a**, Comparison of spike rates for inhibitory vs. excitatory neurons, obtained from ground truth datasets. **b**, Comparison of spike-evoked calcium kernels for inhibitory vs. excitatory neurons, computed as in Fig. 5. Note the different y-axis scaling. **c**, Quantification of the peak of spike-evoked kernels, across neurons. **d**, Comparison of burstiness (coefficient of variation of inter-spike times for inhibitory vs. excitatory neurons. **e**, Example of spike inference from calcium imaging data (top trace, resampled at 30 Hz, at a standardized noise level of “8”, GCaMP8s), with ground truth spike rate (black) as well as spike rates inferred by different variants of the CASCADE algorithm (blue), OASIS (pink) and MLSpike (brown). **f**, Quantification of model performance for ground truth data from inhibitory neurons, resampled at 30 Hz and with a standardized noise level of “8”. **g**, Excerpt from another neuron, highlighting the inability of an algorithm trained with excitatory neurons (GC8-tuned CASCADE) to pick up the non-bursty firing patterns of inhibitory neurons (arrows).

Spike inference with Default CASCADE yielded overall better results, reflecting the model-free approach of CASCADE. CASCADE optimized for the entire GCaMP8 dataset further improved performance, and CASCADE optimized with interneuron ground truth improved even further (p = 0.003 when compared to all model-based approaches; p = 0.03 when compared to Default CASCADE or GC8-tuned CASCADE; repeated-measures ANOVA with Tukey-Kramer correction; Fig. 6e,f). Due to the small interneuron ground truth dataset, we used transfer learning to optimize CASCADE for interneuron data (see Methods). While CASCADE trained on excitatory neuron ground truth (GC8-tuned) was often unable to correctly estimate low-spike-rate episodes without bursts, the same algorithm trained on interneuron ground truth with leave-one-out cross-validation performed noticeably better under these conditions (arrows in Fig. 6g; Fig. 6e). Despite these improvements, spike inference applied to interneurons yielded signals that remained much less correlated with ground truth when compared to other datasets (Fig. Suppl. 1-4).

Together, our analyses highlight the challenges posed by the high spike rates, the small spike-triggered calcium response, and the low burstiness of interneurons for inferring spiking from calcium imaging data. We demonstrate that CASCADE, as a supervised method without prior assumptions about the spike-calcium relationship, is particularly well-suited for this complex task.

## Discussion

The practical value of new tools depends not only on their quality but also on rigorous validation and clear guidance for use. In this study, we examined the calcium indicator GCaMP8, focusing on how existing algorithms for spike inference, which were optimized for previous generations of calcium indicators, need to be adjusted for use with GCaMP8. Through our analysis, we provide specific guidelines, outline important caveats, and offer pretrained models for spike inference with GCaMP8. Together, these insights and resources enhance the interpretability of calcium imaging data, with GCaMP8m and GCaMP8s as promising relatively linear calcium indicators that enable more quantitative spike inference.

The standout feature of GCaMP8 is its fast kinetics. We find that its rise time is sufficiently fast to be particularly beneficial for online spike inference applications such as brain-machine-interfaces, where rapid readout of neuronal activity is critical. Surprisingly, GCaMP8f as the “fast” variant did not exhibit a marked advantage over other GCaMP8 variants for this application. In addition, indicators from previous generations (GCaMP6f; ref. ^8^) achieved rise times almost comparable to GCaMP8 when expressed transgenically, offering an alternative approach that could also be suited for online processing. Overall, our analyses showed limited added benefit from GCaMP8f for population calcium imaging in practice. For special applications, e.g., to study fast calcium dynamics in spines, or to precisely determine inter-spike intervals^35,50^, GCaMP8f may be a highly useful calcium indicator. However, for typical population imaging, we recommend GCaMP8m and GCaMP8s as more practical choices.

GCaMP8f has been recommended for use with fast-spiking interneurons due to its fast kinetics^27^. Our analyses of interneuron data is based on only few interneurons, pooled across different GCaMP8 variants, and we therefore cannot make any conclusive statement about the suitability of different GCaMP8 variants for interneurons. In general, we show that spike inference is less accurate for interneurons than for excitatory neurons (Fig. Suppl. 1-4). Our analyses demonstrate that model-based algorithms designed for cortical pyramidal neurons struggled with interneuron data, while the model-free supervised CASCADE method performed better (Fig. 6). Still, spike inference with interneurons remains challenging, mainly due to the smaller spike-triggered ΔF/F amplitude (Fig. 6b,c) and the reduced bursting tendency (Fig. 6d) of interneurons. The reduced burstiness results in slowly changing calcium signals, from which spike-related calcium increases are more difficult to detect among shot noise compared to the sharper calcium transients typical for excitatory neurons. This interpretation is in line with previous findings from spinal cord neurons, which exhibited lower burstiness similar to cortical interneurons and were more difficult to process for spike inference, in particular with classical model-based methods^42^.

Perhaps the most impactful advancement of GCaMP8, as revealed by our analyses, is its increased linearity. This property has several specific implications for practical applications of spike inference. First, spike rates are inferred more accurately across low and high firing frequencies with GCaMP8 (Supplementary Note 1). Second, high-frequency events are reflected more accurately by ΔF/F due to reduced history-dependent calcium binding dynamics (Fig. 2). Third, the more linear transfer function improves the detection of single action potentials (Fig. 3), particularly for GCaMP8m and GCaMP8s. Furthermore, our analyses demonstrate that calcium indicators of the GCaMP6 family, which cannot detect single action potentials under realistic conditions (Fig. 3) would be able to do so if they were linear but unchanged in terms of kinetics or response magnitudes (Fig. 4). The linearity of GCaMP8m/s in the low-firing regime is therefore highly desirable. However, it also requires spike inference models specifically trained for GCaMP8 to fully benefit from it. We openly provide such pretrained models for spike inference based on the CASCADE framework across a wide range of imaging rates and noise levels (https://github.com/HelmchenLabSoftware/Cascade).

The detection of single action potentials is one of the main goals of calcium imaging. The distinction between single action potentials and bursts is particularly important to understand the nature of the neuronal code in brain areas such as neocortex or hippocampus, where bursts have been implicated in intracellular signaling to control plasticity^38,51^. Earlier indicators like OGB-1 have been shown to enable the detection of single action potentials for selected high-quality recordings^10^. For genetically encoded calcium sensors, detectable calcium transients have been demonstrated to be triggered by single action potentials in rare cases for GCaMP5^26^ and more reliably for selected recordings with YC3.60^12^ or GCaMP6^23^. However, previous studies^8,14,40^ and our work here demonstrate that single action potentials cannot be reliably detected by GCaMP6 variants in cortical pyramidal cells under conditions typical for population imaging (standardized noise levels between 2 and 8; Fig. 3). By contrast, we find that GCaMP8m and GCaMP8s do enable reliable detection of action potentials and therefore re-define the standard. We provide algorithms that are capable of extracting this information (Fig. 3). Interestingly, the earlier indicator XCaMP-Gf (ref. ^25^) also performed well in single action potential detection when noise levels were sufficiently low; however, it has been demonstrated that the necessary brightness and therefore a low noise level is difficult to achieve with XCaMP-Gf using practical laser powers (ref. ^27^), making it a less suitable calcium indicator for most use cases compared to GCaMP8m/s.

A key insight from our analysis is that the linearity of a calcium indicator is an essential factor to enable the detection of individual action potentials (Fig. 4). We believe that this insight will be important for the future engineering of calcium indicators. In this context, it is crucial to understand that the nonlinearity is not only a property of the calcium indicator but also depends on the neuron of interest – ideally, the most linear regime of the indicator (centered around the dissociation constant K_d_) should match the calcium concentration range in which the neuron is physiologically active. In addition, our result on the effect of indicator linearity for the detection of single action potentials will be equally important for experimenters when selecting indicators to resolve small events. From our analyses, we conclude that experiments which require the highest sensitivity (detection of individual somatic spikes, detection of local dendritic events) should be performed with GCaMP8s or GCaMP8m. GCaMP8s, due to its high sensitivity, has been shown to saturate at high spike rates^27^, which may affect its suitability for the recording of neuronal activity, but our analyses suggest that saturation is not strong for cortical principal cells up to spike rates of 15 Hz; the highest overall linearity was, however, achieved with GCaMP8m (Supplementary Note 1). Of course, the ability of GCaMP8m/s to reveal single action potentials will be compromised for high noise levels (Fig. 3) or when neurons cannot be properly distinguished from neuropil background due to dense labeling or low imaging resoluion, and this ability needs to be confirmed also for other brain regions and cell types in the future. Interestingly, detection of single action potentials with GCaMP8f has recently been shown in another brain region in cerebellar granule cells^35^.

One main factor that limits the accuracy of spike inference is the unexplained variability of calcium responses across cells and cell types. For example, the transfer functions for GCaMP8s demonstrate that the inferred spike rate is approximately linear for individual neurons but that the slope of this linearity is different from neuron to neuron. This variability across neurons has been identified before^42,43^ and could be due to different calcium concentration levels^43^ or different calcium buffering conditions across neurons^52,53^, or due to systematic errors when computing ΔF/F_0_, in particular when working with the low baseline levels (F_0_) of GCaMP indicators^42^. Here, we extracted the unitary response to single action potentials from calcium imaging data, with the hope that scaling the inferred spike rates with this factor would collapse these individual transfer functions, a procedure known as auto-calibration^15,43^. However, although single action potentials could be inferred more reliably after auto-calibration (Fig. 3e-g), the same procedure had negative effects for the regime of higher spike rates, most likely due to the remaining nonlinearity of GCaMP8 (Fig. Suppl. 3-4). In our opinion, this analysis highlights the two most important requirements towards fully accurate calcium imaging: the development of even more linear calcium indicators, and the development of algorithms to perform auto-calibration while taking into account the residual nonlinear behavior. Here, we demonstrate that our algorithms for spike inference together with GCaMP8m/s are close to achieving this goal. Together, our analysis is therefore both a step towards more interpretable spike inference and a potential steppingstone for the next advance of quantitative spike inference.

## Contributions

P.R. conceived the project, analyzed data, and wrote the initial draft. M.R. collected the GCaMP8 data that were re-analyzed for this manuscript and contributed ideas for data analysis. X.F. performed the auto-calibration analysis and contributed to the writing of the manuscript. K.S. supervised data collection. F.H. guided the project and contributed to the writing of the manuscript.

## Acknowledgements

This work was supported by a grant from the Swiss National Science Foundation (Ambizione grant PZ00P3_209114 to P.R.).

## Data availability

New ground truth data for GCaMP8, GCaMP7f and X-GCaMP-Gf, including extracted spike times and calcium traces, are deposited in the GitHub repository together with demo scripts (https://github.com/HelmchenLabSoftware/Cascade). We provide a cloud-based Colaboratory Notebook that allows for interactive browsing through all datasets (https://colab.research.google.com/github/HelmchenLabSoftware/Cascade/blob/master/Demo%20scripts/Explore_ground_truth_datasets.ipynb). Other already existing and publicly available datasets are described in detail in the Methods (‘Ground truth datasets and quality control’).

## Code availability

A cloud-based version of CASCADE that includes pretrained deep learning-based models for GCaMP8 variants and pretrained models for online spike inference is available as a Colaboratory Notebook (https://colab.research.google.com/github/HelmchenLabSoftware/Cascade/blob/master/Demo%20scripts/Calibrated_spike_inference_with_Cascade.ipynb). The code is also available as a GitHub repository together with demo scripts, installation instructions and FAQs (https://github.com/HelmchenLabSoftware/Cascade). Pretrained models for CASCADE are archived in an online server (https://www.switch.ch/drive/) and retrieved automatically by the CASCADE code.

## Methods

### Ground truth datasets and quality control

Datasets comprising simultaneous fluorescence imaging of cellular calcium signals and electrophysiological juxtacellular recordings of spikes from the same neurons (“ground truth” datasets) were retrieved from publicly available databases and quality-controlled for each neuron as described previously^14^. Specifically, ground truth datasets for virally induced GCaMP6 expression were obtained from the primary publication related to the introduction of GCaMP6 (ref. ^23^). Ground truth datasets for transgenically expressed GCaMP6 were obtained for mice from a dedicated publication from the Allen Institute^41^ and from the CASCADE publication for zebrafish^14^. All these datasets and associated quality controls have been described previously^14^.

For ground truth dataset recorded with the three GCaMP8 variants, as well as ground truth datasets recorded with GCaMP7f and XCaMP-Gf, we used the openly accessible data provided with the primary publications introducing GCaMP8 (ref. ^27,28^). We performed additional quality controls as we did for previous similar analyses^14^ and excluded ground truth recordings from neurons that exhibited excessive motion artifacts, too high spike rates (indicative of unidentified interneurons), or that did not allow the accurate isolation of electrophysiological spikes from the recorded neuron. These cleaned-up datasets for XCaMP-Gf, GCaMP7f, GCaMP8f, GCaMP8m and GCaMP8s are available in addition to previously uploaded datasets in a format that can be accessed both in MATLAB and Python (datasets DS#28-DS#32 on https://github.com/HelmchenLabSoftware/Cascade).

### Standardized noise levels and resampling of ground truth data

Standardized noise levels were obtained^14,42^ by computing the median absolute fluctuation of *ΔF/F* between adjacent timepoints, normalized by the square root of the imaging frame rate (*f*_*r*_). This metric quantifies shot noise in *ΔF/F*, which is independent across consecutive time points and can therefore be estimated from their pairwise differences. The median provides a robust measure of the typical fluctuation while reducing the influence by outliers caused by fluorescence transients upon action potentials.

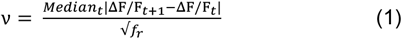

When computed from ΔF/F data, the standardized noise level *ν* is normalized by the frame rate *f*_*r*_ and therefore quantitatively comparable across datasets even when frame rates differ, hence the name “standardized noise levels”. The units for *ν* are %·Hz^−1/2^, which we omit in the main text for readability.

To obtain ground truth for well-defined and comparable conditions, we used the same procedures as previously described^14^. Briefly, fluorescence data from ground truth datasets were first temporally resampled to the desired sampling rate using the *signal*.*resample()* function from the Python package SciPy^54^. Then, to obtain a target noise level *ν*, Gaussian noise was added until the noise metric *ν* yielded the desired level. As a side effect, Gaussian white noise affects all temporal frequencies equally and is thus unbiased with respect to indicators exhibiting different rise and decay times. We have previously shown that such addition of Gaussian noise is, for this purpose, a sufficient approximation of typical Poisson-distributed shot noise^14^. In practice, this shot noise reflects the influence of many separate factors, such as the level of calcium indicator expression, the laser power used for excitation, the quantum efficiency of the specific indicator, the photodetection efficiency, and the number of pixels within a neuronal ROI.

### OASIS for spike inference

For spike inference using Default OASIS, we employed the Python implementation provided in Suite2p^31^ (https://github.com/MouseLand/), running in Python 3.7 with default parameters. To identify the optimal value of the time constant – the main free parameter of the algorithm – we performed a systematic grid search across a wide range of possible values (Fig. Suppl. 1-3). This procedure yielded a dataset-specific, optimized version referred to as *Fine-tuned OASIS*. For each dataset, the time constant that maximized the correlation between inferred and ground truth spike rates (median across neurons) was selected and subsequently applied to all neurons in that dataset. The optimal values for the time constant parameter, optimized for ground truth at a standardized noise level of 8, were 1.4 s for AAV-GCaMP6f, 1.6 s for AAV-GCaMP6s, 0.45 s for TG-GCaMP6f, 0.95 s for TG-GCaMP6s, 0.45 s for GCaMP7f, 0.2 s for GCaMP8f, 0.45 s for GCaMP8m, 0.5 s for GCaMP8s, and 0.001 s for GCaMP8-INH.

### MLSpike for spike inference

The MLSpike algorithm was downloaded from https://github.com/MLspike/spikes and used within MATLAB 2022b (ref. ^55^). To set up Default MLSpike (that is, a version of MLSpike that does not explicitly require ground truth), the inverse frame rate *dt* as a parameter of the algorithm was fixed to the value constrained by the ground truth dataset. The *drift* parameter was set to 0.1. The nonlinearity parameter was set to 0.1. The Hill coefficient was set to 1.84 for GCaMP6s, 2.99 for GCaMP6f, 3.1 for GCaMP7f, and 2.08/1.92/2.2 for GCaMP8f/m/s, as reported in previous publications. The parameters *amplitude* (amplitude of a single action potential in ΔF/F, in %), *tau2* (rise time constant, in milliseconds) and *tau* (decay time constant, in milliseconds) were set to 0.113, 0.034, 0.121, 0.294, 0.511 and 0.576 for *amplitude*, 70.2, 15.6, 15.3, 2.96, 3.31 and 4.72 for *tau2*, and 1870, 760, 129, 57, 107, and 267 for *tau* (for GCaMP6s, GCaMP6f, GCaMP7f, GCaMP8f, GCaMP8m and GCaMP8s). This version of MLSpike is considered “Default” because it takes the physiological values from the relevant papers for GCaMP6 (ref. ^55^) and GCaMP7-8 (ref. ^27^).

To optimize MLSpike for a specific dataset (Fine-tuned MLSpike), the same initial parameters as above were used, but a 2D grid search as described for the OASIS algorithm was performed to obtain the optimal parameters for *amplitude* and *tau* (Fig. Suppl. 1-2). Grid search was performed for each dataset for a standardized noise level of 8, and the best-performing parameters as quantified by correlation with ground truth spike rates were applied to all neurons of this dataset. Due to long processing times with MLSpike, no out-of-dataset generalization was performed. Furthermore, a parameter search in higher-dimensional space, for example by including rise times or nonlinearity parameters, might result in even better performance for MLSpike but is prohibitive due to the slow processing speed of MLSpike. Optimal values obtained for *tau*/*amplitude* were 0.5/2.5 for GCaMP7f, 0.3/2.0 for GCaMP8f, 0.3/1.5 for GCaMP8m, 0.7/1.0 for GCaMP8s, and 2.5/0.2 for GCaMP8-INH.

### Spike inference with CASCADE

For spike inference with Default CASCADE, pretrained models together with the algorithm were downloaded from https://github.com/HelmchenLabSoftware/Cascade. These models had been trained on a large database of excitatory neurons across different brain areas with a focus on cortical recordings using the GCaMP6 indicator (named “global CASCADE models” in ref. ^14^). Here, these models are called Default CASCADE because they were not retrained using GCaMP8 ground truth.

To optimize CASCADE for GCaMP8 ground truth data, several approaches were tested. First, CASCADE was retrained from scratch on all GCaMP8 data (covering all variants, “f”, “m” and “s”). These models are called GC8-tuned CASCADE in this study. Second, CASCADE was retrained from scratch on a specific dataset, e.g., GCaMP8f only. These models are called, e.g., GC8f-tuned CASCADE or Fine-tuned CASCADE (akin to Fine-tuned MLSpike mentioned above). Third, for interneurons, CASCADE was initially trained from scratch on the same datasets as the GC8-tuned CASCADE models (INH-tuned CASCADE), but then the model was retrained with a transfer-learning approach^56^ with dataset-specific ground truth while freezing the weights for all but the last layer (INH-transfer CASCADE). This procedure benefits from convolutional filter weights obtained by fitting a large and diverse ground truth and fine-tuning the weighting of these filters for the specific ground truth dataset. Similar transfer learning procedures have been used successfully in neuroscience applications to reduce the amount of required training data^57,58^.

In all instances of retraining, CASCADE networks consisted of a standard convolutional network with six hidden layers, including three convolutional layers. The input consisted of a window of 64 time points (32 time points for frame rates <15 Hz), symmetric around the time point for which the inference was made. The three convolutional layers had relatively large filter sizes (31, 19 and 5 time points; 17, 9 and 3 time points for frame rates <15 Hz), with an increasing number of features (20, 30 and 40 filters per layer), with max pooling layers after the second and third layer, and a densely connected hidden layer consisting of ten neurons as the final layer. To avoid any effect of overfitting on our results, the same neuron was never used both for training and testing of a model. For example, if a model was tested for GCaMP8f data, separate models were trained for each test neuron of the GCaMP8f data while excluding during training the neuron tested for (“leave-one-out” strategy). This cross-validation strategy prevents fitting of test data and enables us to test the generalization of the algorithm to unseen data.

Models retrained with GCaMP8 in general or for specific GCaMP8 datasets at various sampling rates and for a broad range of noise levels are integrated into CASCADE (https://github.com/HelmchenLabSoftware/Cascade); further models tailored towards special use cases can be readily requested as described in the FAQ of the GitHub page.

### CASCADE for online spike inference

For the investigation of online spike inference, specific new CASCADE models were trained for each dataset and each “integration time”. Integration time is here defined as the number of data points in the future from the current time point of inference that can be seen by the spike inference algorithm (Fig. 5c). For example, standard spike inference with CASCADE uses a symmetrical window of 64 time points, with 32 before and 32 after the time point of inference. For online spike inference, the number of time points after the time point of inference was decreased and the number of time points before was increased to maintain a 64-point window. Pretrained CASCADE models for spike inference for GCaMP6 and GCaMP8 for frame rates of 30 Hz and 60 Hz are integrated into CASCADE and are available online: https://github.com/HelmchenLabSoftware/Cascade.

### Benchmarking and comparisons across algorithms

As a potential conflict of interest, some of the authors have been the lead authors of the development of the algorithm CASCADE^14^. To avoid a bias towards this algorithm during benchmarking comparisons (Fig. 1 and Fig. 6), we did not construct new evaluation metrics but used existing ones designed and used previously (Pearson’s correlation between smoothed spike trains) ^17,18,14^. For other algorithms (MLSpike and OASIS), we used grid search to find optimal parameters. Grid search, at least for MLSpike, is computationally expensive but ensures unbiased optimization, in contrast to gradient-based optimization, which is faster but might be used in a suboptimal manner without being obvious. Furthermore, we did not apply the leave-one-neuron-out cross-validation strategy for grid search with MLSpike and OASIS (for reasons of computational resources required for cross-validation), in contrast to training with CASCADE, where we employed this strategy. The parameters found for MLSpike and OASIS should therefore be considered a “fit” of the data, not a cross-validated model as for CASCADE. This difference in procedure should, if anything, give an advantage to MLSpike or OASIS compared to CASCADE. To quantify spike inference performance, ground truth spike rates used for training and evaluation were generated from discrete ground truth spikes by convolution with a Gaussian smoothing kernel. The precision of the ground truth was adjusted by tuning the standard deviation of the Gaussian smoothing to the temporal sampling rate (σ = 0.05 s for 30 Hz recordings and σ = 0.025 s for 60 Hz recordings). Next, this smoothed ground truth spike rate was compared to the inferred spike rate using Pearson’s correlation. For evaluation of spike rate inference results with OASIS and MLSpike, we tested Gaussian smoothing kernels of variable standard deviation and temporal shifts between -1 and +1 s to find the amount of smoothing and the delay for each dataset that optimized the correlation with ground truth spike rates. Additional metrics were used for specific analyses as described in the main text.

### Analysis of deviations from linearity and from identity

To compute the deviation from linearity (Supplementary Note 1), the mean normalized squared deviations of the inferred spike rate, *SR*_*inferred*_, from the linear fit to the inferred spike rate, *SR*_*linear*_, were computed. These deviations were averaged across spike rate bins of the transfer function (bins *k* from 1 to N in the equation below) and then converted to regular normalized units by taking the square root. This metric corresponds to the mean square root error but is normalized to obtain relative deviations from linearity.

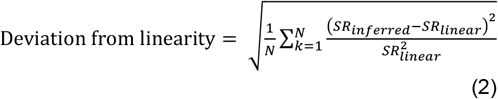

To compute the deviation from identity for the transfer function of true spike rate to inferred spike rate (Supplementary Note 1), a linear fit *y* = *m* · *x* was computed, with the true spike rate *x*, the inferred spike rate *y*, and the slope of the linear transfer function fit, *m*. The deviation from identity was defined as:

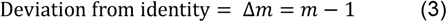

### Analysis of high-frequency events

For detection of high-frequency events, ground truth spiking binned to 30 Hz was smoothed with a 35-point running box filter. Then, all contiguous time windows with the smoothed ground truth spike rate >6 Hz were labeled using the MATLAB function *regionprops()* and defined as events. For such events, the time bin of each supra-threshold time window with the highest spike rate (“peak time”) was retrieved. 50 time points before and 50 points after peak time were extracted as a local excerpt for this event. The goal of this procedure was to define events where a relatively large number of spikes occurred within a short time window.

The number of such extracted events across neurons were, with dataset ID in brackets: 92 (DS#9, GC6f, virally induced), 329 (DS#10, GC6f, transgenic), 281 (DS#11, GC6f, transgenic), 174 (DS#13, GC6s, transgenic), 36 (DS#14, GC6s, virally induced), 265 (DS#29, GC7f), 712 (DS#30, GC8f), 1070 (DS#31, GC8m) and 1218 (DS#32, GC8s). Dataset IDs follow the ground truth database at https://github.com/HelmchenLabSoftware/Cascade.

### Extraction of linear response kernels from ground truth data

Linear kernels of the calcium signal in response to an average action potential were extracted by regularized deconvolution using the *deconvreg*(*Calcium,Spikes*) function in MATLAB (MathWorks). These linear kernels are shown in Fig. 5a and Fig. 6b. This function computes the kernel, which, when convolved with the observed *Spikes*, results in the best linear approximation of the *Calcium* trace. These kernels are used to simulate linear calcium traces while otherwise keeping response magnitudes and kinetics intact for the analyses in Fig. 4a-e.

### Simulation of nonlinear calcium dynamics

To simulate nonlinear responses to single action potentials *de novo* (Fig. 4f), we implemented a Hill nonlinearity^9^, which models cooperative binding of proteins and is used to describe the nonlinear relationship between calcium concentrations and the resulting fluorescence:

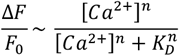

For the purpose of the analyses, we kept n = 2 constant and varied the inflection point of the nonlinearity, K_D_. In this scenario, a lower K_D_ results in higher linearity in the low-firing regime, and vice versa. This implementation of nonlinearity is only a simple approximation of the more complex nonlinearities and history-dependencies found in reality^16,35^ but serves as a model to highlight and dissect the effect of AP-evoked signal strength (which is a product of indicator brightness and dynamic range) and indicator linearity.

To simulate linear and nonlinear calcium dynamics for Supplementary Note 1, we followed existing work that models the relationship of spikes and fluorescence in linear and nonlinear manner^6^. Briefly, we defined a kernel with infinitesimal rise time, a decay time of 200 ms and a single-spike response amplitude of 10% ΔF/F. Action potential times were added at manually chosen time points for illustrative purpose and convolved with the single-spike response kernel. The nonlinear fluorescence 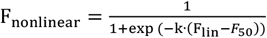 was obtained from the linear fluorescence F_lin_ with parameters F_50_ = 65% and k = 0.2. The resulting nonlinear trace was rescaled by a linear fit to match the overall scale of the linear fluorescence trace F_lin_.

### Quantal analysis using Gaussian mixture models for auto-calibration

For auto-calibration (Fig. 3e-g), minimal calcium transients were detected and assumed to be related to isolated single (“unitary”) action potentials. To robustly detect minimal events, a fine-tuned version of CASCADE (e.g., GC8s-tuned CASCADE for GCaMP8s data) was applied to ΔF/F traces to obtain a denoised inferred spike rate. Contiguous periods of the inferred spike rate that were above the expected threshold θ of a single spike were defined as events (e.g., θ = 0.3 as the half of the maximum spike rate to be expected from a single inferred spike for a CASCADE model trained with a ground-truth smoothing standard deviation of 50 ms and a sampling rate of 30 Hz). The inferred spike rate within each of these events was summed, resulting in a histogram of summed spike rates across events for each neuron. A modified Gaussian mixture model (GMM) in MATLAB was then used to perform a quantal analysis from the histogram to infer the unitary amplitude. To account for prior knowledge about the distribution of event amplitudes and to improve the GMM fit, the initial amplitude of unitary events, *A*, was initialized to expected amplitudes (*A* = 1.6). Furthermore, as is common for quantal analyses, the fitting procedure took into account that Gaussian components were expected to have integer relationships, with the distribution having peaks at events with 1, 2, 3 and 4 discrete spikes. To this end, a hierarchical structure of the GMM fit was established by initializing means as integer multiples of the expected unitary amplitude *A*. Additionally, the variance of each component was assumed to follow a variance scaling parameter that grew proportionally with the mean of each component. The histogram of event-related spike rates was then fit to extract the unitary amplitude. The inferred unitary amplitude was then used to rescale the entire inferred spike rates. For events as defined above which contained exactly a single ground truth spike, the spike rate inferred with CASCADE and the spike rate inferred with CASCADE corrected by the unitary amplitude factor were compared to the true spike number (Fig. 3g). We provide a MATLAB toolbox to perform auto-calibration using quantal analysis of inferred spike rates from CASCADE via GitHub: https://github.com/PTRRupprecht/Quantal-analysis-for-spike-inference.

### Statistical analysis and box plots

Statistical analyses were performed in MATLAB 2022b. The Mann–Whitney rank-sum test was used for non-paired samples, and the Wilcoxon signed-rank test was used for paired samples. Two-sided tests were applied unless otherwise stated. ANOVA with Tukey-Kramer correction was used when multiple comparisons were considered. Box plots used standard settings in MATLAB, with the central line at the median of the distribution, the box at the 25th and 75th percentiles and the whiskers at extreme values excluding outliers (outliers defined as data points that are more than 1.5·D away from the 25th or 75th percentile value, with D being the distance between the 25th and 75th percentiles).

## Supplementary Figures

**Figure 1 Supplement 1.**
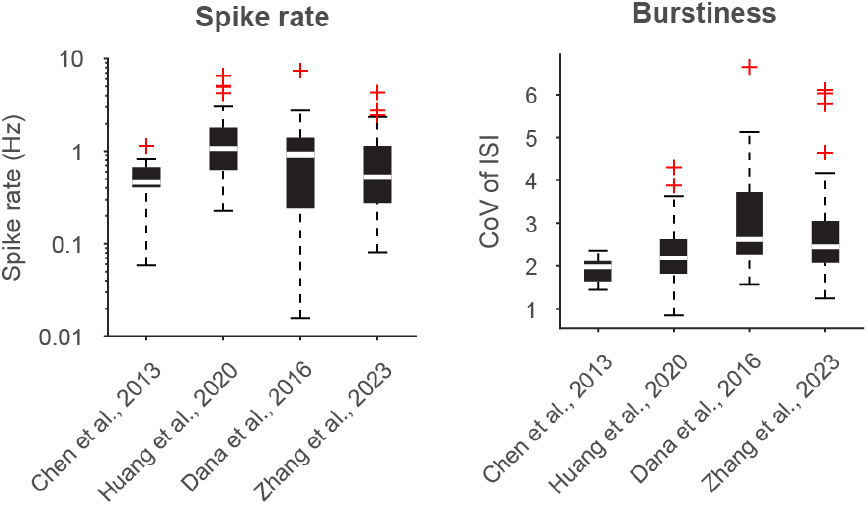
Electrophysiological properties of GCaMP8 ground truth in comparison to existing ground truth recordings. Comparison of ground truth recordings for GCaMP6 (virally induced) (ref. ^23^, Chen et al.), GCaMP6 (transgenic expression) (ref. ^8^, Huang et al.), jRGECO1a/jRCaMP1a (ref. ^29^, Dana et al.), as well as GCaMP8 (ref. ^27^, Zhang et al.), all recorded in excitatory L2/3 neurons from mouse visual cortex during anesthesia. **Left panel:** Spike rates of recordings from Zhang et al. (0.53 [0.28; 1.14] Hz, median [inter-quartile range], n = 137 neurons) were not statistically different (p > 0.05, Wilcoxon rank-sum test) from the spike rates from Chen et al. (0.46 [0.41; 0.67] Hz, n = 18) and Dana et al. (0.91 [0.24; 1.39] Hz, n = 20), and lower than the spike rates from Huang et al. (1.07 [0.63; 1.82] Hz, n = 74; p < 0.0001). Therefore, spike rates from Zhang et al. are in the same range as previous recordings. **Right panel:** Burstiness was quantified as the coefficient of variation (CoV) of inter-spike intervals (standard deviation of inter-spike intervals divided by the mean spike interval). CoV values of recordings from Zhang et al. (2.4 [2.1; 3.0]) were slightly but statistically significantly higher than for the recordings by Chen et al. (2.0 [1.6; 2.1], p = 0.03, ANOVA with Tukey-Kramer correction) but not statistically different from the recordings by Huang et al. and Dana et al. (2.2 [1.8; 2.6], p = 0.52 and 2.62 [2.3; 3.7], p = 0.33). Therefore, recordings by Zhang et al. are slightly more bursty than most other ground truth recordings from visual cortex.

**Figure 1 Supplement 2.**
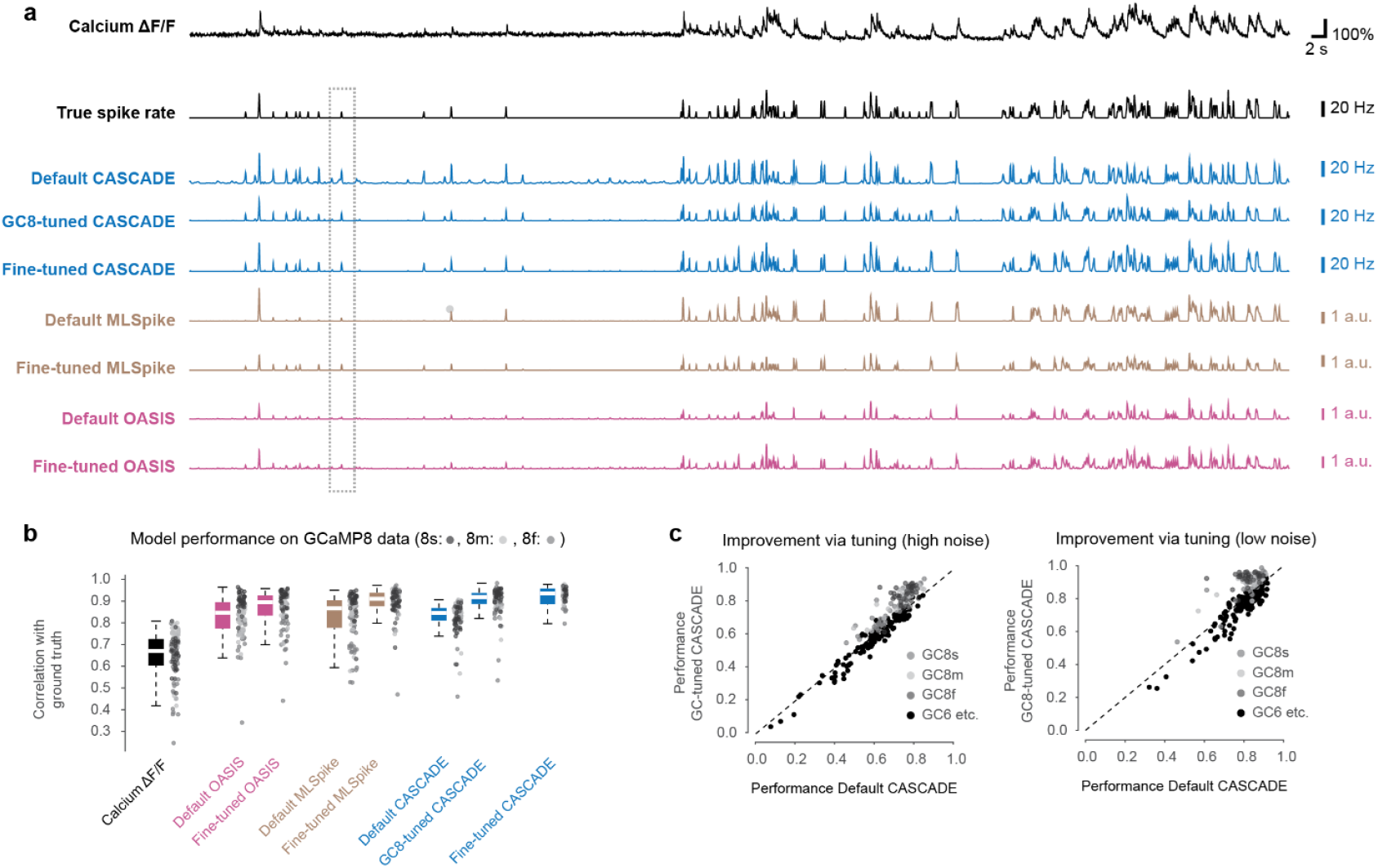
Benchmarking of spike inference algorithms for GCaMP8 for low noise levels. **a**, Example of spike inference from calcium imaging data (top trace, resampled at 30 Hz, at a standardized noise level of “2”, GCaMP8m), with ground truth spike rate (black) as well as spike rates inferred by different variants of the CASCADE algorithm (blue), OASIS (pink) and MLSpike (brown). GC8-tuned models were trained with the ground truth available across all GCaMP8 variants, while fine-tuned models were trained based only on the variant of interest (e.g., GCaMP8m). Spike inference for the same recording but resampled at higher noise levels is shown in Fig. 1d. The dashed box highlights the example of an isolated action potential, shown also in Fig. 1f. **b**, Quantification of model performance across all GCaMP8 ground truth data resampled at 30 Hz and with standardized noise level of “2”. **c**, Performance of spike inference (correlation with ground truth) for Default CASCADE (trained with pre-GCaMP8 data) vs. GCaMP8-tuned CASCADE. Spike inference for GCaMP8 data is improved, performance for data from other ground truth datasets, primarily GCaMP6, is decreased, for both high and low noise levels.

**Figure 1 Supplement 3.**
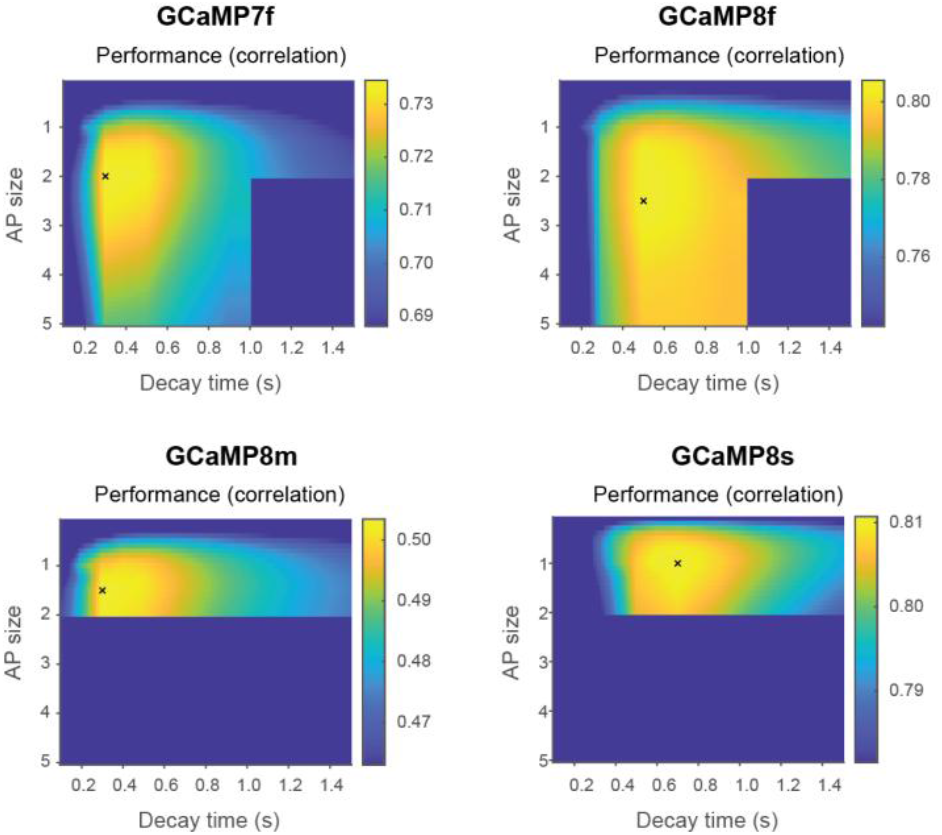
Grid search for optimization of MLSpike parameters for spike inference with GCaMP8. Search was performed in a 2D parameter space spanned by the two parameters *amplitude (AP size)* and *tau* (*decay time)*. Please note the different scaling of the four color maps; further, notice that the range of values is relatively narrow for all color maps and does not go to zero performance (typically spanning a range of <10% of the average value). Due to the high computational cost of running MLSpike, parameter search was stopped when the most likely global maximum was found, resulting in empty values of the grid search (dark blue).

**Figure 1 Supplement 4.**
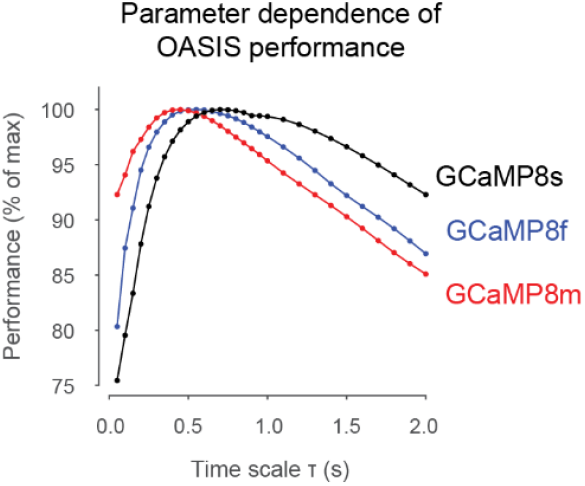
Grid search for optimization of OASIS for spike inference with GCaMP8. Search was performed in a 1D parameter space (*time scale*). Note that the y-range of values is relatively narrow and does not cover performance <75% of the maximum.

**Figure Supplement 5.**
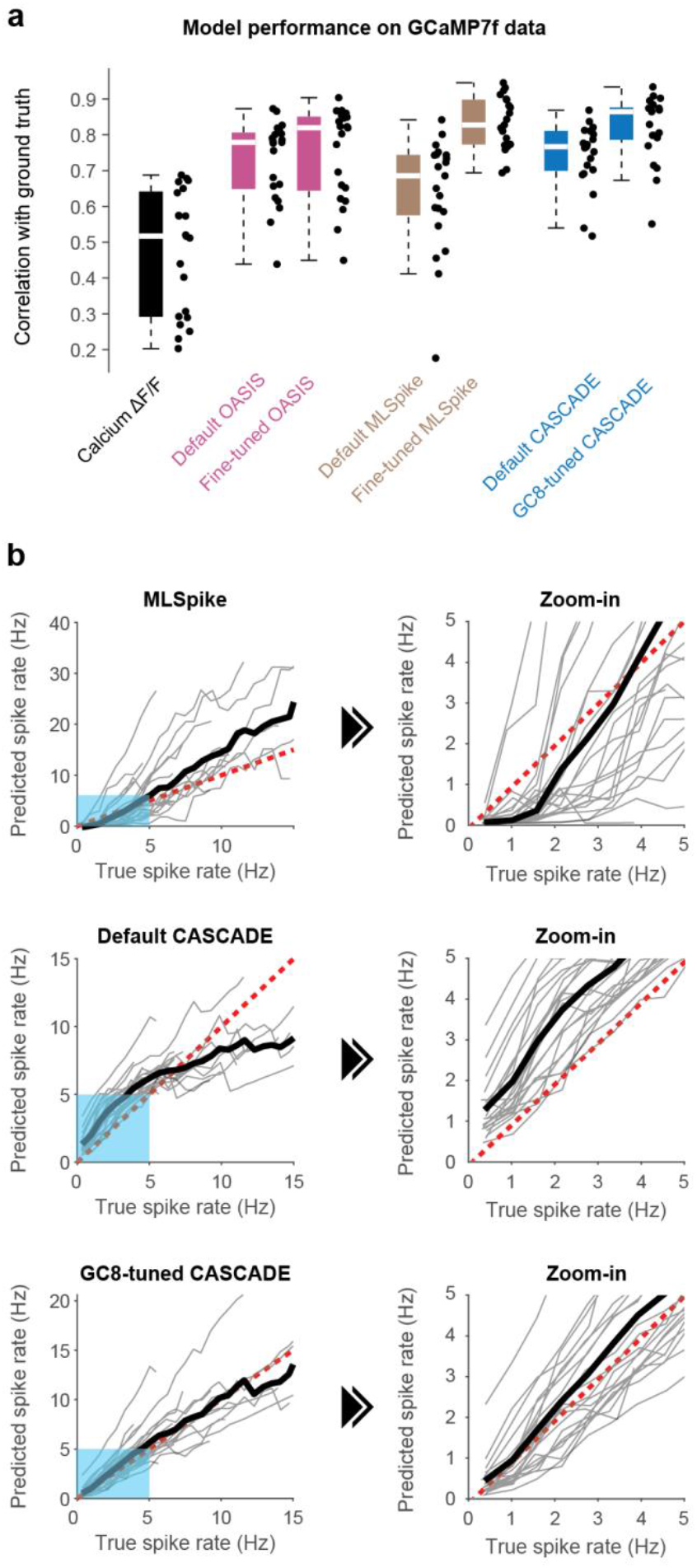
Benchmarking of spike inference algorithms for GCaMP7f. **a**, Quantification of model performance (correlation with ground truth) across all GCaMP7f ground truth data resampled at 30 Hz and with standardized noise level of “8”. The best-performing algorithm was CASCADE trained with GCaMP8 ground truth (0.86 [0.79; 0.88], median [IQR] across 21 neurons), which was significantly better than Default CASCADE (0.77 [0.7; 0.81]; p = 1.1e-5, repeated-measures ANOVA with Tukey-Kramer correction), although not significantly different from the fine-tuned versions of OASIS (p = 0.10; 0.82 [0.64; 0.85]) and MLSpike (p = 0.99, 0.83 [0.77; 0.9]). **b**, Transfer function to visualize the average spike rate as obtained with different spike inference algorithms as a function of true spike rate for the GCaMP7f dataset, as in Supplementary Note 1. The dashed red line represents the unity relationship. MLSpike underestimates absolute spike rates for low-frequency firing rates and results in overestimates for high frequencies. The Default CASCADE mode overestimates low firing frequencies and underestimates high firing frequencies, an effect already seen before^14^. The model trained on GCaMP8 data yielded relatively unbiased estimates both for low and high firing rate regimes. All analyses are limited by the small GCaMP7f dataset (21 neurons) and should ideally be confirmed for additional datasets recorded with GCaMP7f.

**Figure 1 Supplement 6..**
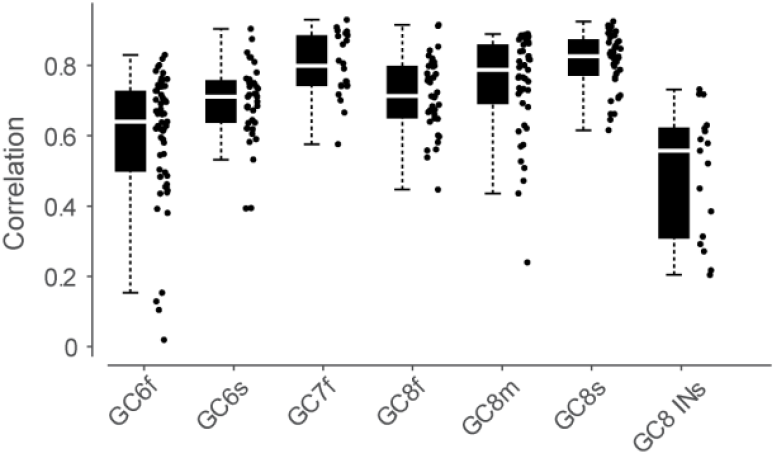
Performance across datasets. A CASCADE model was trained on a specific dataset and tested on the same dataset, using the leave-one-neuron-out strategy for cross-validation. The resulting spike inference performance (correlation) can in theory be used as an indicator for the quality of a calcium indicator. This quantification, however, needs to be interpreted very carefully since the datasets were not necessarily recorded by the same experimenter and using the same depth of anesthesia or sensory stimuli (cf. also Fig. Suppl. 1-1). For example, higher technical recording quality would result in higher correlation, similar to more frequent or stronger evoked stimulation due to the increase of signal compared to noise in the recording.

**Figure 3 Supplement 1.**
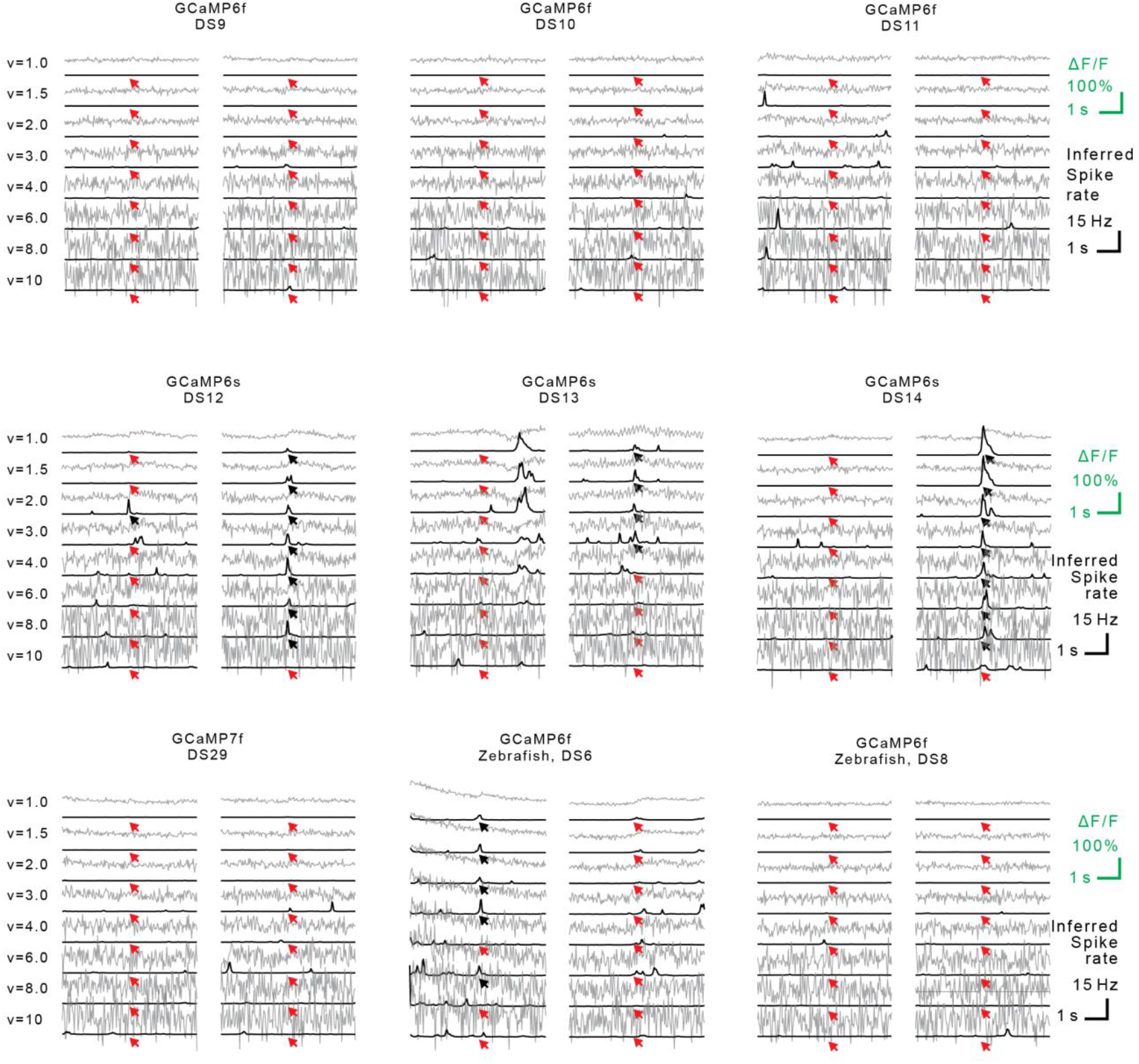
Examples of calcium signals triggered by single action potentials. Examples of isolated action potentials with the corresponding ΔF/F trace (gray) and the inferred spike rate (black) for an increasing standardized noise level ν. Arrows indicate the time point when the true isolated action potential occurred (red arrow if not detected). Examples shown for different datasets with GCaMP6 variants and with GCaMP7f, with two examples from different neurons for each dataset shown in the two columns (see Methods for details about the datasets).

**Figure 3 Supplement 2.**
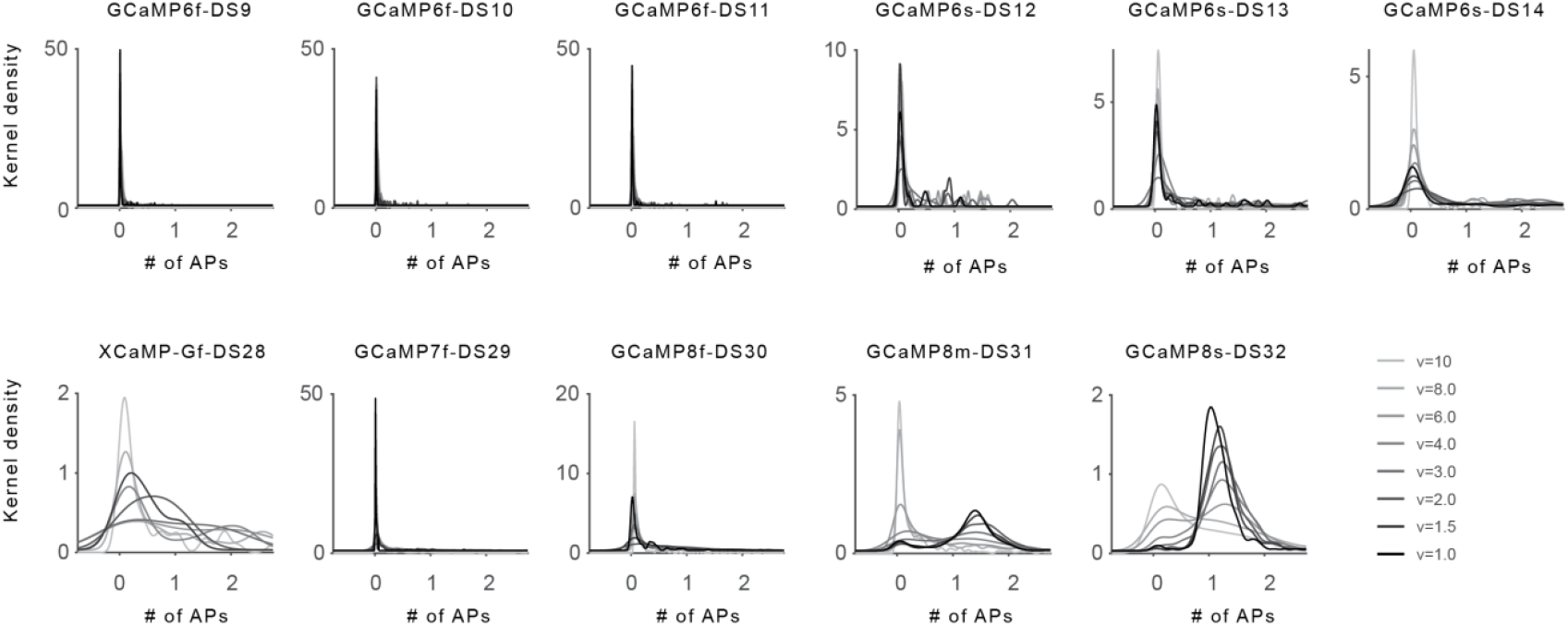
Distributions of inferred spike numbers for single isolated action potentials. Extension of Fig. 3b. Number of detected action potentials (APs) for a true isolated single AP, plotted as a distribution (kernel density estimate). Shades of gray indicate the different noise levels (lowest noise level, dark grey; highest noise level, light grey, as indicated in the legend at the bottom right).

**Figure 3 Supplement 3.**
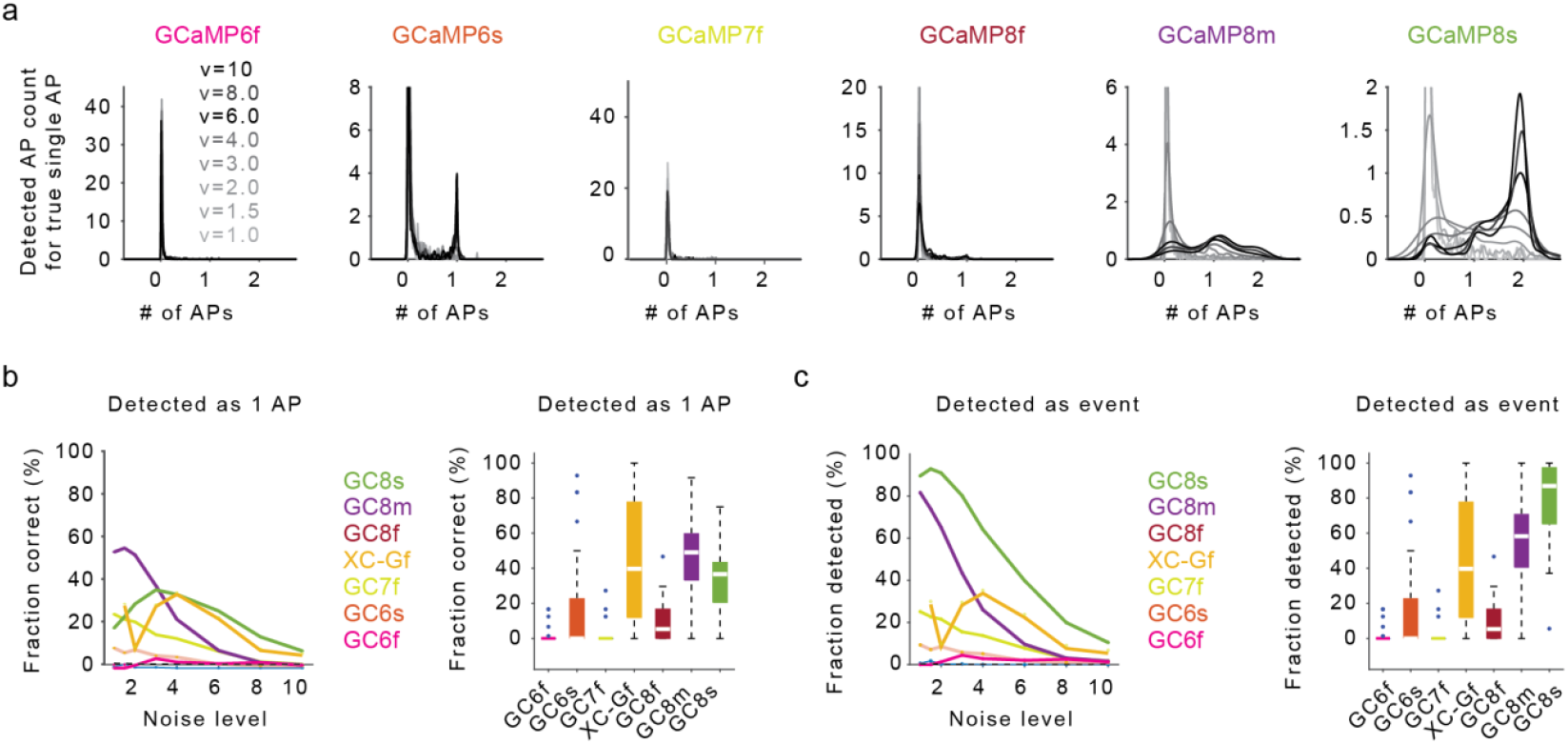
Detection of isolated action potentials from GCaMP8 with spike inference using MLSpike. This figure qualitatively replicates the findings from Fig. 3b-d but using MLSpike instead of CASCADE as an algorithm for the detection of action potentials. Based on these data, the overall performance of MLSpike for detection of single action potentials is overall slightly worse compared to CASCADE. However, the comparisons across indicators exhibit the same qualitative differences and trends as shown in Fig. 3. **a**, Number of detected action potentials (APs) for a true isolated single AP, plotted as a distribution (kernel density estimate). Shades of gray indicate the different noise levels (lowest noise level, dark grey; highest noise level, light grey). Ideally, the distribution would be narrow and centered around “1” true action potential. **b**, *Left:* Fraction of APs correctly detected as a single AP (spike count >0.5 and <1.5) across datasets and noise levels. *Right*: Fraction of APs per neuron correctly detected as a single AP, averaged across noise levels ν = 2-4 for robustness. **c**, *Left:* Fraction of APs correctly distinguished from noise (spike count >0.5) across datasets and noise levels. *Right*: Fraction of APs correctly distinguished from noise, averaged across noise levels ν = 2-4 for robustness.

**Figure 3 Supplement 4.**
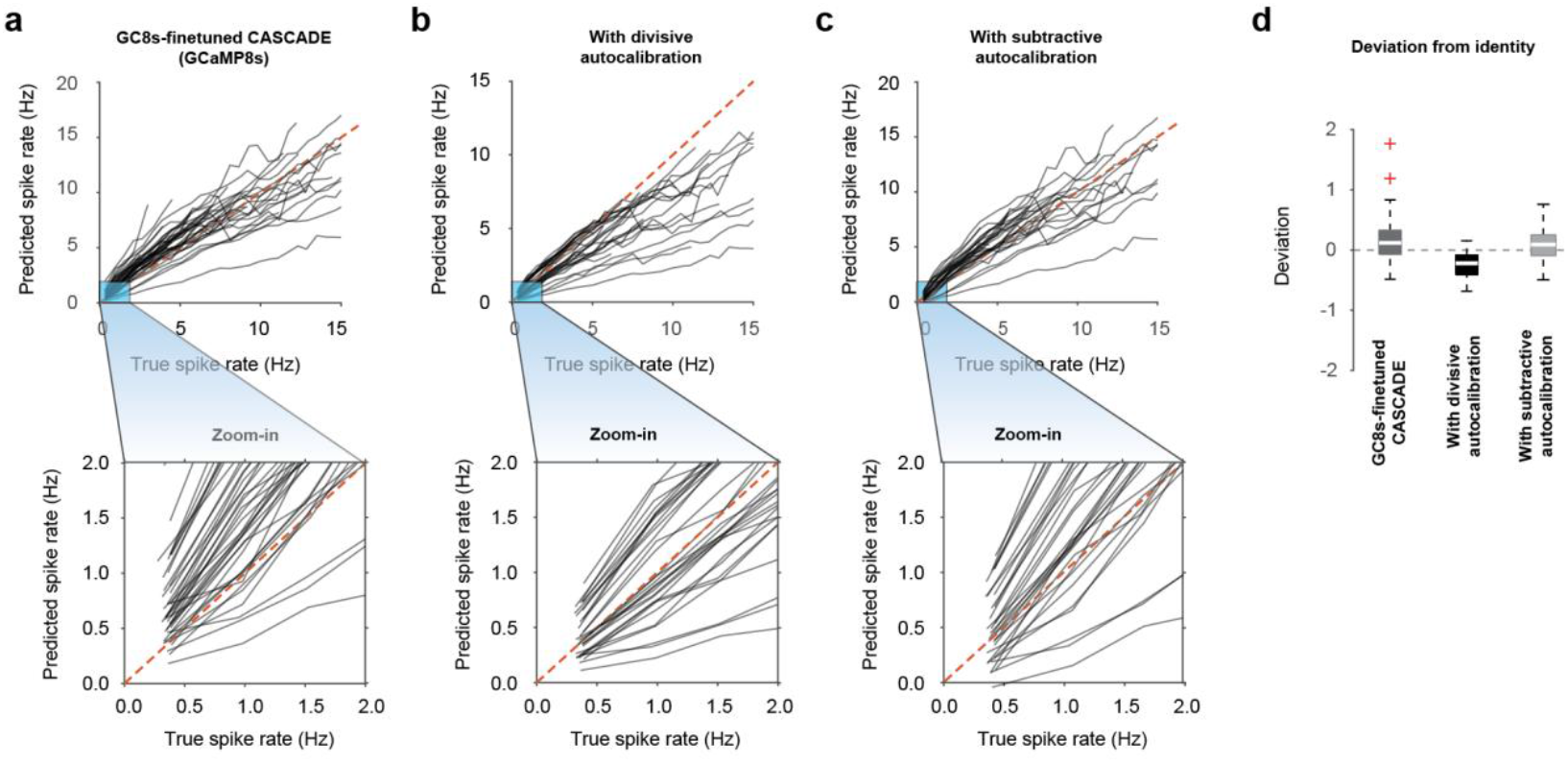
Effect of auto-calibration on the deviation from identity. **a-c**,The transfer functions (panels a-c) are displayed as in Fig. 2. Auto-calibration is performed by either dividing the inferred spike rate by the unitary scaling factor (“divisive auto-calibration”, panel b) or by subtracting the excess of unitary response globally (“subtractive auto-calibration”, panel c). After divisive auto-calibration (cf. Fig. 3h-i), the spike rates inferred for events with low spike rates are less biased (zoom-in; compare to the red identity line). However, the inferred spike rates for high true spike rates are quenched, resulting in a deviation from identity that is, in absolute terms, not significantly improved compared to the spike rates before auto-calibration (panel d; p = 0.40, Wilcoxon signed-rank test). After subtractive auto-calibration, the spike rates inferred for events with low spike rates are also less biased (zoom-in) but exhibit a higher dispersion around the identity line for the lowest spike rates compared to divisive auto-calibration (zoom-in). The inferred spike rates for high true spike rates are largely unchanged, resulting in a deviation from identity that exhibits a trend to be slightly lower than for the spike rates before auto-calibration (panel d; p = 0.08).

**Figure 5-1.**
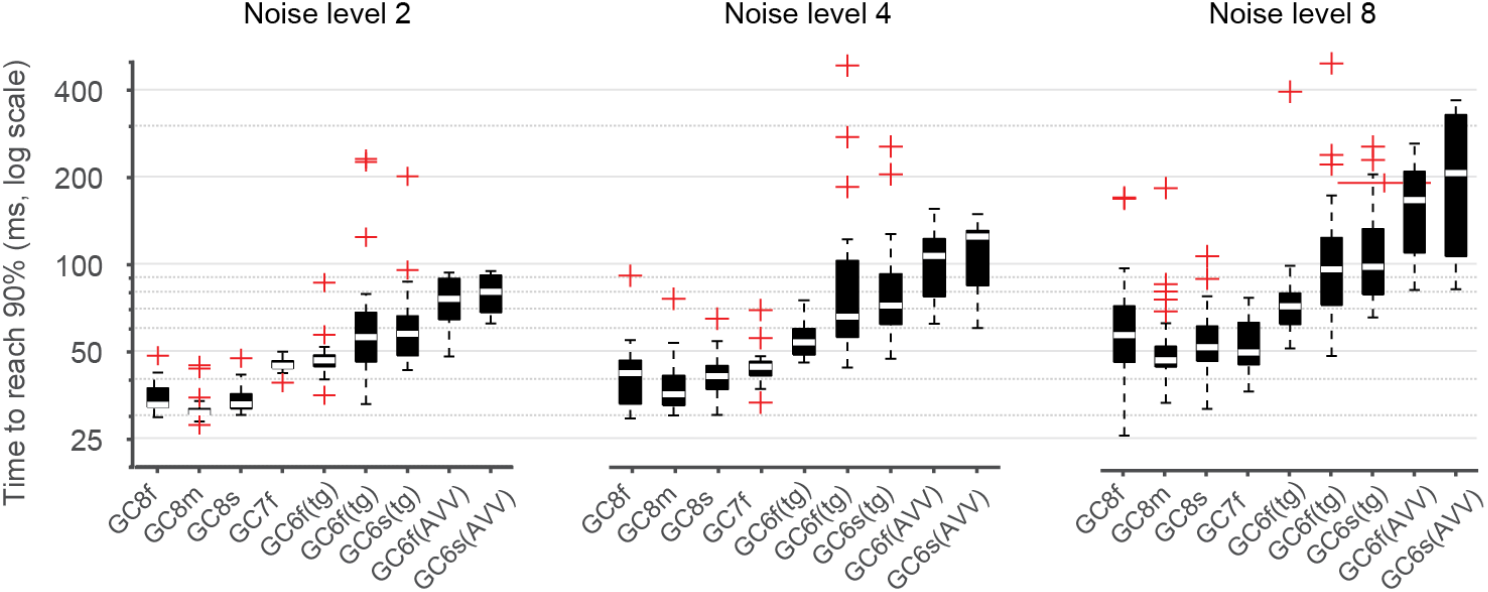
Online spike inference performance across noise levels. Same quantification as in Fig. 5e but for variable amounts of standardized noise levels. The minimal delay achieved with GCaMP8 indicators rises with noise levels from ∼30 ms for noise level 2 to >50 ms for noise level 8.

## Supplementary Table

**Suppl.Table S1.**
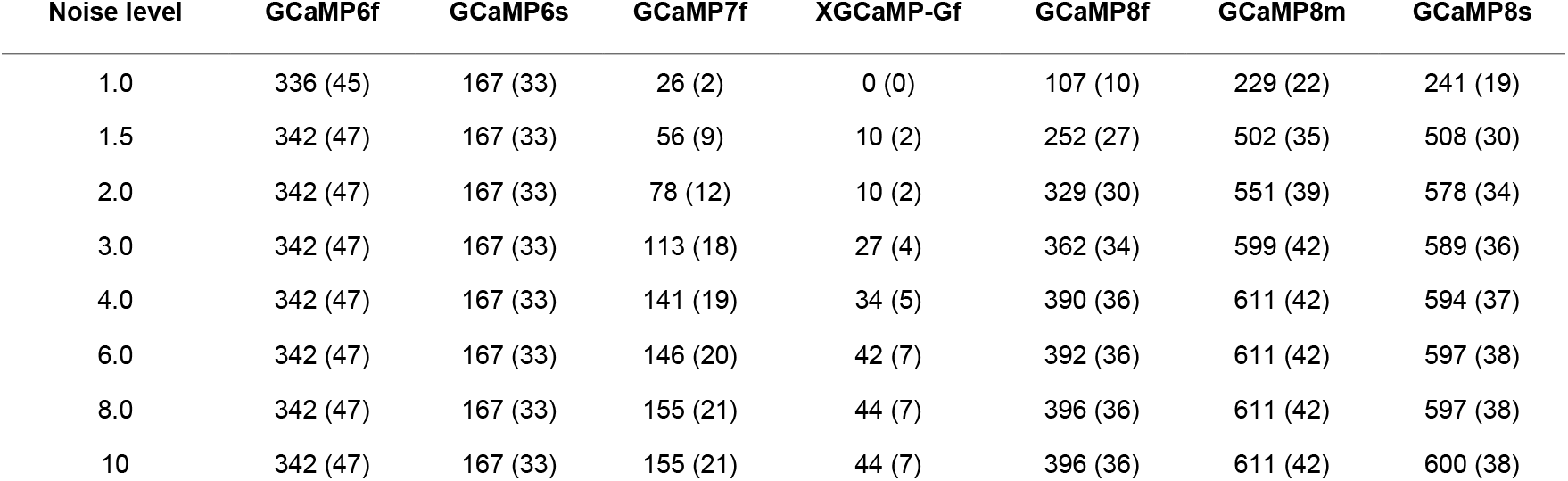
Number of isolated action potentials detected. For the analyses in Fig. 4, Gaussian noise was added to ground truth recordings to obtain data with a defined noise level (left-most column). Since some ground truth recordings were already performed at a relatively high noise level, they could not be included for low-noise condition analyses. Therefore, the number of included isolated action potentials (values indicated in the dataset-specific columns) and the number of the underlying neurons (values indicated in brackets) increased with analyses of higher noise levels.

| Noise level | GCaMP6f | GCaMP6s | GCaMP7f | XGCaMP-Gf | GCaMP8f | GCaMP8m | GCaMP8s |
| --- | --- | --- | --- | --- | --- | --- | --- |
| 1.0 | 336 (45) | 167 (33) | 26 (2) | 0 (0) | 107 (10) | 229 (22) | 241 (19) |
| 1.5 | 342 (47) | 167 (33) | 56 (9) | 10 (2) | 252 (27) | 502 (35) | 508 (30) |
| 2.0 | 342 (47) | 167 (33) | 78 (12) | 10 (2) | 329 (30) | 551 (39) | 578 (34) |
| 3.0 | 342 (47) | 167 (33) | 113 (18) | 27 (4) | 362 (34) | 599 (42) | 589 (36) |
| 4.0 | 342 (47) | 167 (33) | 141 (19) | 34 (5) | 390 (36) | 611 (42) | 594 (37) |
| 6.0 | 342 (47) | 167 (33) | 146 (20) | 42 (7) | 392 (36) | 611 (42) | 597 (38) |
| 8.0 | 342 (47) | 167 (33) | 155 (21) | 44 (7) | 396 (36) | 611 (42) | 597 (38) |
| 10 | 342 (47) | 167 (33) | 155 (21) | 44 (7) | 396 (36) | 611 (42) | 600 (38) |

## Supplementary Note 1: Estimation of absolute spike rates across firing frequencies

Ideally, spike inference would provide not only an estimate proportional to spike rate but also reveal the correctly scaled absolute spike rate. However, existing algorithms for spike inference typically struggle to achieve this when applied to datasets that are not part of the training data^14^. Moreover, they often do not even attempt to estimate absolute spike rates and instead rely on “correlation with ground truth” as a scaling-invariant metric of performance^18^. CASCADE was designed to infer absolute spike rates for unseen datasets and, as a state-of-the-art algorithm, managed to reveal absolute spikes with an error factor of ∼2 for indicators like GCaMP6f/s (ref. ^14^). This means that estimated spike rates typically fall within 50% to 200% of the true spike rate^14^. In this section, we investigate how well absolute spike rates can be estimated across different neuronal firing rates from GCaMP8 data.

To visualize predicted vs. true spike rates, we smoothed both ground truth and predicted spike rates with a temporal Gaussian filter (standard deviation, 1 s) and binned the resulting ground truth data into firing frequency bins with increments of 0.6 Hz. This enabled us to plot ΔF/F and predicted spike rates as a function of the true spike rate for each neuron, creating a “transfer function” (Fig. N1a-e). A striking feature of these transfer functions, shown for the GCaMP8s dataset, is the nonlinearity observed with Default CASCADE (Fig. N1b). This property has previously been noted for CASCADE when trained with GCaMP6 data (Fig. 4g in ref.^14^). In contrast, our analysis demonstrates that GCaMP8 does not require this trade-off, as CASCADE variants trained with GCaMP8 ground truth can recover a more linear transfer function (Fig. N1c-e).

**Figure for Supplementary Note 1 (Fig. N1).**
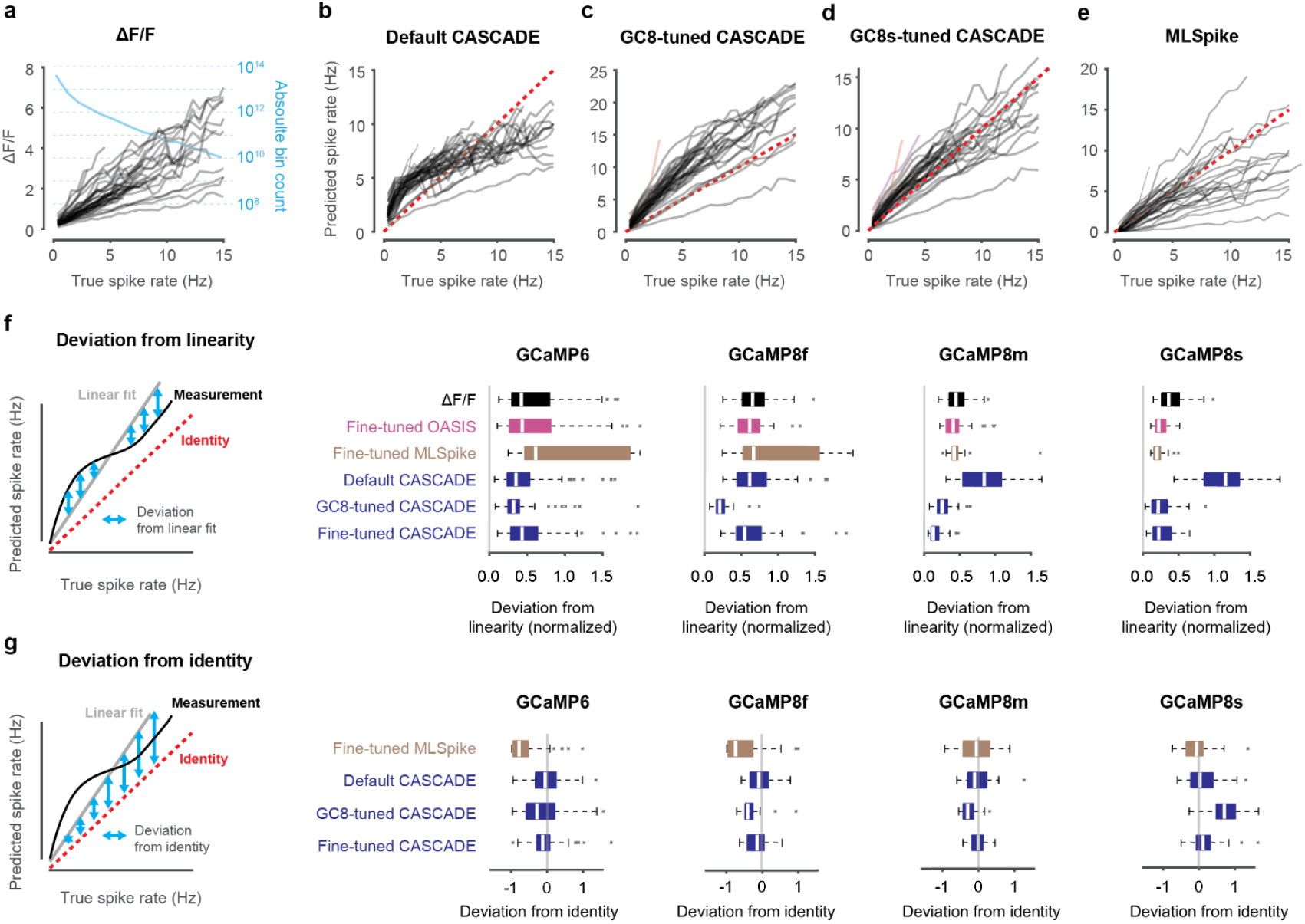
Inference of absolute spike rates: Deviations from linearity and identity. **a**, Average ΔF/F as a function of true spike rate for the GCaMP8s dataset. Each connected line represents the values from a single neuron. The blue line indicates the frequency of each true spike rate on a log scale. For example, 15 Hz events were almost 10^4^ times less likely to occur than silence. **b-e**, Average spike rate as obtained with Default CASCADE (b), GC8-tuned CASCADE (c), GC8s-tuned CASCADE (d) and fine-tuned MLSpike (e), as a function of true spike rate for the GCaMP8s dataset. The red dashed line represents the unity relationship. **f**, Left: Schematic of assessing the deviation (blue arrows) of the experimental data for each neuron (black curve) from linearity (gray line). Right panels: Normalized deviations from linearity for different indicators and algorithms across neurons. **g**, Left: Schematic of assessing the deviation (blue arrows) of the linear fit to the experimental data (gray line) from identity (red dashed line). Right panels: Deviation from identity for different indicators and algorithms across neurons.

To systematically quantify the linearity of the transfer function across datasets, we calculated the “deviation from linearity” for each neuron and algorithm (Fig. N1f). This metric measures the relative amount of variation that is not captured by a linear fit and therefore contributes to the nonlinearity (see Methods for definition). These findings shed light not only on the nonlinearities of the different indicators but also how these are dealt with by the different algorithms. OASIS, which is based on a linear model of calcium dynamics^59^, did not introduce additional nonlinearities compared to raw ΔF/F. In contrast, MLSpike increased the nonlinearity for some indicators but not others. Interestingly, the deviation from nonlinearity was relatively low for Default CASCADE when applied to GCaMP6 (0.35 [0.23; 0.53]) and to GCaMP8f (0.62 [0.44; 0.84]), but high for GCaMP8m and GCaMP8s datasets (0.84 [0.54; 1.09] and 1.16 [0.86; 1.35]). These findings indicate that the nonlinear behavior of GCaMP6 is captured by Default CASCADE and transfers relatively well to GCaMP8f but not to GCaMP8m/s. However, when retrained with GCaMP8 data, CASCADE exhibited a substantial and consistent decrease of nonlinearity compared to ΔF/F when applied to GCaMP8 data. Overall, when tested on all GCaMP8 data, GC8-tuned CASCADE resulted in significantly more linear predictions than all other algorithms (p < 10^-5^ for all algorithms except for Fine-tuned CASCADE, where p = 0.02; repeated-measures ANOVA with Tukey-Kramer correction; median deviation from linearity with inter-quartile ranges: 0.49 [0.32; 0.66] for ΔF/F, 0.38 [0.25; 0.55] for OASIS, 0.42 [0.24;0.55] for MLSpike, 0.85 [0.55; 1.20] for Default CASCADE, 0.22 [0.16; 0.31] for GC8-tuned CASCADE, and 0.26 [0.13; 0.48] for fine-tuned CASCADE). These results demonstrate that supervised models like Default CASCADE incorporate the nonlinearities present in their training data (GCaMP6) and need to be retrained (GC8-tuned CASCADE) for less nonlinear indicators (GCaMP8) to optimize the linearity of spike inference.

We reasoned that the higher linearity of the GCaMP8 datasets would also lead to CASCADE models that behave more linearly in general. To test the linearity of spike inference models, we applied CASCADE models to synthetic ground truth that was simulated with a linear kernel and was based on the spike patterns from electrophysiological ground truth recordings^23,29,8,27^. We found that CASCADE models trained with GCaMP8 data behaved much more linearly compared to models trained with previous indicators (Fig. N2).

**Figure for Supplementary Note 2 (Fig. N2).**
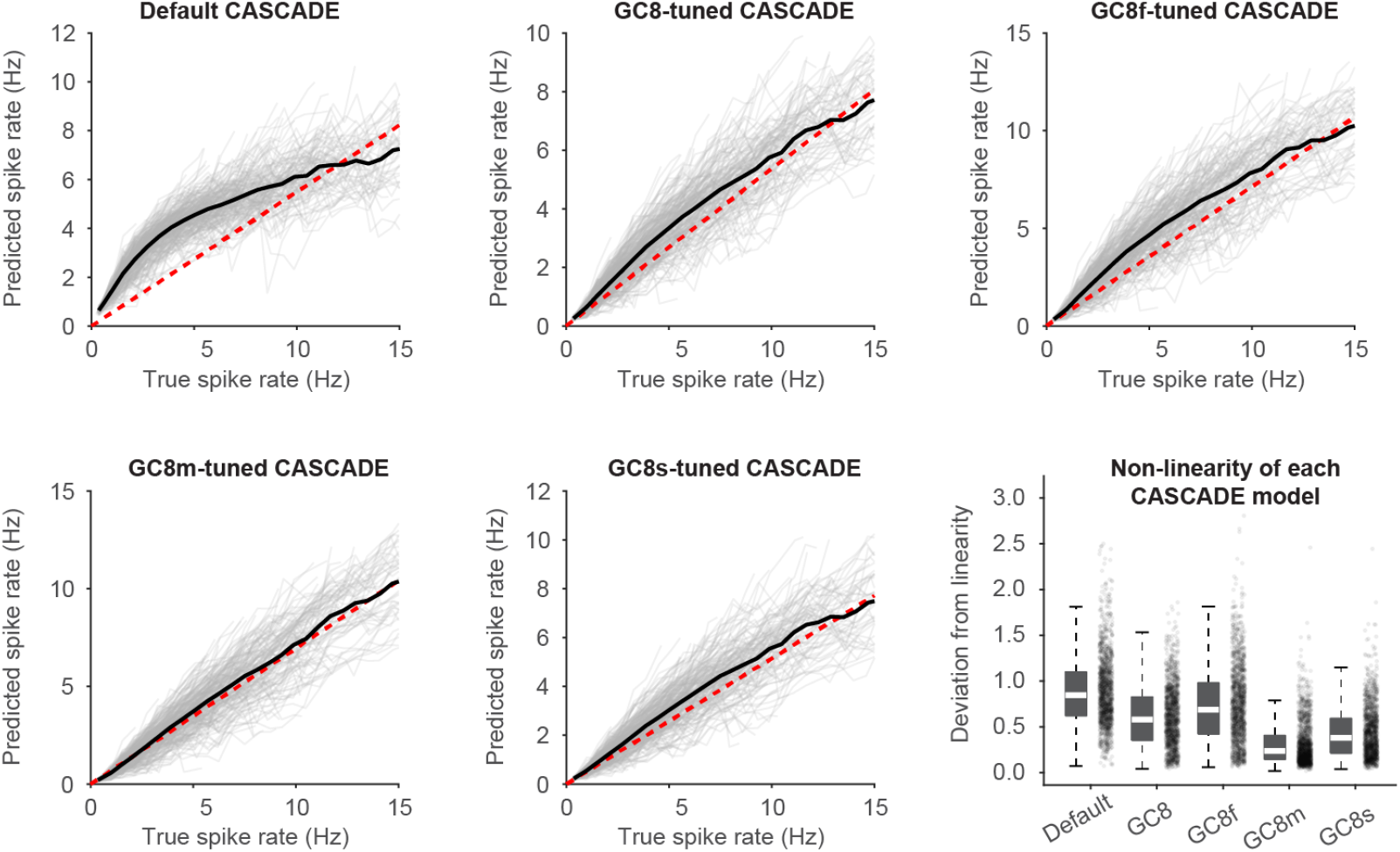
Evaluation of nonlinearities using a synthetic ground truth. To evaluate the transfer function of various CASCADE models and their nonlinearity, a synthetic ground truth was generated. The ground truth spike patterns consisted of experimentally recorded action potentials of excitatory neurons from refs. ^8,23,27,29^ (n = 237 neurons with variable recording duration, each represented by a gray line; 102,093 action potentials, 32.7 hours of recordings) to ensure naturalistic spike patterns. These experimentally obtained spike patterns were convolved with a double exponential kernel (rise time 5 ms, decay time 500 ms, peak amplitude 40% ΔF/F), with Gaussian noise added to reach a standardized noise level of “8”. Spike inference algorithms were applied to this entire linear synthetic dataset. From the inferred spike rates and the synthetic ground truth, transfer functions as in Fig. N1 were retrieved, allowing to judge the linearity of the models trained on specific datasets. The Default CASCADE model exhibited the highest nonlinearity (black average transfer curve) compared to the weighted linear fit of the inferred spike rates (red dashed line); the sublinearity of the average transfer function at high firing rates reflects the supralinearity of the training data. CASCADE trained with GCaMP8 ground truth yielded a relatively linear transfer function, with mild but clear signs of saturation for models fine-tuned for GCaMP8f and GCaMP8s. The transfer function obtained from a model trained with GCaMP8m (GC8m-tuned CASCADE) data was the most linear. Since the models reflect the nonlinearities of their training data, these analyses indicate that GCaMP8, and in particular GCaMP8m, are distinctly more linear calcium indicators than for example GCaMP6 (which is the basis of most of the training data for Default CASCADE) across the firing rate regime that is typically covered by experimentally obtained ground truth recordings. The bottom right panel shows quantified deviations from linearity as introduced in Fig. N1 for the different CASCADE models. Quantifications were pooled across multiple simulations with variable setting of the kernel parameters to emulate GCaMP8f, GCaMP8m, GCaMP8s, GCaMP7f and GCaMP6 with the kernel rise times (2, 3, 4, 11 and 50 ms, respectively), decay times (45, 80, 190, 100 and 300 ms) and peak amplitudes (55, 71, 81, 70 and 25 % ΔF/F). Deviations from linearity were reduced compared to Default CASCADE by 30% (GC8-trained), 19% (GC8f-trained), 60% (GC8m-trained) and 48% (GC8s-trained). Therefore, the GC8m-trained model exhibited the lowest deviation from linearity, suggesting the highest linearity of the GC8m indicator used for its training. All comparisons p < 10^-6^, repeated-measures ANOVA with Tukey-Kramer correction, compared across n = 1,422 instances of simulated neurons.

These results demonstrate that the calcium indicators GCaMP8s and in particular GCaMP8m, and the models trained with these data behave much more linearly than previous indicators and supervised models.

Next, we observed that the transfer function of individual neurons not only deviated from a linear relationship but also from the identity relationship (Fig. N1g). Deviations from the identity line indicate a systematic bias to under- or overestimate spike rates. To quantify the deviation from identity, we calculated Δ*m* as the difference between the slope of the linear fit, *m*, and unity (see Methods; Fig. 2g). A positive value indicates an overestimation of spike rates, while a negative value indicates an underestimation. For this quantification, we excluded raw ΔF/F and the OASIS algorithm, as both do not estimate absolute spike rates.

Our results highlight a key difference between CASCADE models that were trained with all GCaMP8 datasets and those trained on specific GCaMP8 variants, such as GCaMP8s (Fig. N1g). When evaluated across all GCaMP8 data, the models trained with specific ground truth made relatively accurate predictions of spike rates (median Δm across neurons with inter-quartile range 0.20 [0.08; 0.36]) that were more precise than for other algorithms (Δm: 0.44 [0.24; 0.74] for MLSpike, 0.28 [0.16; 0.43] for Default CASCADE, and 0.40 [0.26; 0.59] for GC8-tuned CASCADE; p < 2 10^-4^; repeated-measures ANOVA with Tukey-Kramer correction). Specifially, models trained on all GCaMP8 ground truth (GC8-tuned CASCADE) underestimated spike rates for GCaMP8f and GCaMP8m (Δm = -0.39 [-0.46; -0.24] and -0.30 [-0.43; -0.13]) and overestimated spike rates for GCaMP8s (+0.71 [0.48; 1.02]). Such a bias is not necessarily prohibitive because a spike rate estimate with an uncertainty factor of <2 is still very useful. For example, a Δm of +0.71 indicates a systematic bias to overestimate spike rates by ∼71%, which seems acceptable for many applications. This bias for the GCaMP8-tuned model needs to be weighed against the larger and more diverse dataset used to train it, which has been shown to be advantageous for the robustness of an algorithm^14^. However, if the goal of the analysis is to estimate absolute spike rates as accurately as possible, our analyses suggest that training a CASCADE model on the specific GCaMP8 variant is beneficial.

